# Genome-wide association study in European beech (*Fagus sylvatica* L.) for drought stress traits

**DOI:** 10.1101/2025.04.08.647712

**Authors:** Mila Tost, Ourania Grigoriadou-Zormpa, Selina Wilhelmi, Timothy Beissinger, Mehdi Ben Targem, Markus Müller, Henning Wildhagen, Alexandru Lucian Curtu, Oliver Gailing

## Abstract

Forest tree breeding is an extremely long and tedious process. To study the genetic architecture of polygenic traits in long-lived species such as forest trees, costly field experiments are implemented. Phenotypic data on traits, that are measured at maturity, are only available after a long time and juvenile-mature correlations are often unknown. Genome-wide association studies (GWAS) aim to identify loci associated with drought stress, tree physiology, growth or wood quality traits, which could be prioritized in breeding programs. Genotypic and phenotypic data were collected from approximately 100 adult beech trees per stand in five locations in the South-Eastern Romanian Carpathians along an altitudinal gradient associated with precipitation and temperature. We performed GWAS using PLINK to identify SNP markers associated with traits related to drought stress. A total of 121 markers on eight chromosomes were identified as being associated with stomatal density. Sixty-four markers are located on chromosome 10 in a region spreading from ∼4.89 to 13.67 Mb. There are five genes in this region that are thought to play a role in controlling stomatal density. All markers within this region exhibit similar allele frequencies, which are correlated with stomatal density and the altitudinal gradient of the stands. We assume this entire region is jointly involved in local adaptation to drought stress. We identified one interesting candidate SNP associated with leaf nitrogen content. Two SNP markers were identified as being significantly associated with δ^13^C as measure of intrinsic water use efficiency. Additionally, signals of significant polygenic selection for δ^13^C were observed.

## Introduction

In forest tree breeding, trees from different populations exposed to specific environmental conditions are sampled and tested in provenance trials or common garden experiments (Neale and Kremer 2011). Adult trees are later tested, because growth and quality trait measurements from juvenile trees are not reliable (Neale and Kremer 2011). In the face of climate change, provenance trials are also used to identify drought tolerant trees with desirable wood quality and growth traits for forest tree breeding programs (Neale and Kremer 2011; Rellstab et al. 2015; Cortés et al. 2020). This step already takes between 20 to 25 years in long-living woody perennials like European beech (*Fagus sylvatica* L.) (Neale and Kremer 2011) and only then the seed and seedling production starts (Neale and Kremer 2011; Cortés et al. 2020). Therefore, it is of great interest in forest tree breeding to determine the genetic basis of adaptive and economic traits and the potential for success of selection on a trait, before we start selection (Cortés et al. 2020; Degen and Müller 2023). Furthermore, identifying candidate gene markers associated with specific traits is desired, as they could be utilized in breeding programs (Neale and Kremer 2011; Cortés et al. 2020; Müller et al. 2024).

Genome-wide association studies (GWAS) aim to identify associations between markers and phenotypic traits of interest (Visscher et al. 2017; Neale and Kremer 2011). GWAS are becoming more popular in forest genetics research (Cumbie et al. 2011; Cappa et al. 2013; Diao et al. 2024; Meger et al. 2024) with increasing availability of reference genomes for tree species (Nystedt et al. 2013; Mishra et al. 2022). A limited number of GWAS have been conducted in European beech (Pfenninger et al. 2020; Müller et al. 2024; Lazic et al. 2024), and even less GWAS used the chromosome-level assembly of European beech (Lazic et al. 2024; Pfenninger et al. 2021).

GWAS return peaks of the “genomic landscape” of adaptation (Rellstab et al. 2015), but according to the “infinitesimal model” of Fisher (1918), quantitative traits are controlled by many small-effect loci. Several methods have been introduced based on Fisher’s infinitesimal model (Berg and Coop 2014; Racimo et al. 2018; Zeng et al. 2018, Josephs et al. 2019). Another approach using the “infinitesimal model” is referred to as Ĝ (Beissinger et al. 2018). Ĝ follows the model of Wright (1931) that under a selective pressure, alleles with positive effect sizes increase in frequency over generations. In the Ĝ test, all loci are considered together to retrieve a signal of polygenic selection (Beissinger et al. 2018). Ĝ is calculated by estimating marker effects with RR-BLUP (Endelmann 2011), an estimation approach from genomic prediction (Beissinger et al. 2018; Mahmoud et al. 2023). But instead of predicting phenotypes or breeding values, Ĝ uses the estimated marker effects and combines them with allele frequency differences calculated between different groups that have encountered a selective pressure (Beissinger et al. 2018; Mahmoud et al. 2023). Ĝ offers an advantage over genomic prediction by providing information on the adaptive or selective pressures that have shaped a trait, and can even indicate the direction of selection. This allows us to determine whether a trait has been subject to negative or positive selection (Beissinger et al. 2018; Mahmoud et al. 2023). Ĝ can be used to identify signals of polygenic selection when effect sizes are too small to pass the significance threshold of a GWAS or other methods like selection signature mapping or environmental association analysis (Beissinger et al. 2018; Mahmoud et al. 2023). Furthermore, Ĝ has the advantage over genomic prediction in that its results are easier to interpret (Mahmoud et al. 2023). Ĝ does not identify individual markers associated with phenotypes (Beissinger et al. 2018; Mahmoud et al. 2023), but the estimated marker effects can be used for prediction of the performance of individuals (Meuwissen et al. 2001; Endelmann 2011; Alves et al. 2020). Therefore, Ĝ is a promising new approach to test for polygenic selection under an environmental selective pressure (Beissinger et al. 2018). Studies in crop species have shown that Ĝ successfully identifies signals of polygenic selection (Morales et al. 2021; Mahmoud et al. 2023).

Our study comprises 514 individual trees collected in the South-Eastern Carpathian Mountains (Romania) from five European beech stands, along an altitudinal gradient which follows a precipitation gradient. The mean annual precipitation was higher at the higher altitudes; hence the lower stands are more likely to experience periods of limited soil water availability. Several drought stress related and physiological traits were measured. These traits include δ^13^C in leaf dry matter as a proxy for water use efficiency (Fotelli et al. 2003; Querejeta et al. 2003) and stomatal density. Stomata, located in the plant leaf epidermis, are important to control gas exchange and transpiration (Gailing et al. 2008; Bertolino et al. 2019; Chen et al. 2022). Stomata characteristics as stomatal density and control are important in managing changing CO_2_ levels, temperatures, photosynthetically active radiation and water availability (Gailing et al. 2008; Bertolino et al. 2019; Chen et al. 2022). Furthermore, the leaf nitrogen content and the leaf carbon content were measured, and based on these measurements the C/N ratio was calculated. Leaf nitrogen content and the C/N ratio are impacted by complex physiological mechanisms (Gessler et al. 2017). At the start of a drought event, lower nitrogen uptake is observed, which later leads indirectly to carbon starvation, which further reduces nitrogen uptake (Gessler et al. 2017). Drought stress can lead to short-term depletion of leaf nitrogen and long-term depletion of leaf carbon (Gessler et al. 2017; Ruehr et al. 2019). The impact on the C/N ratio depends on the nutrient availability before the drought stress event, lower nutrient availability at the beginning of drought stress will result in a higher C/N ratio during drought (Gessler et al. 2017). The correlation of minor allele frequencies (MAF) observed at GWAS candidate loci and traits of interest were further investigated. Nearby annotated genes were referenced based on a literature review to evaluate their putative role in local adaptation.

The findings from this GWAS could help us to prioritize candidate SNPs for drought stress traits in forest breeding programs. If traits are too polygenic to identify candidate SNPs, Ĝ can provide information on whether traits have a genetic basis and were under polygenic selection. When Ĝ is successful, the previously estimated marker effects could also be used for genomic prediction. To our knowledge, Ĝ has not been applied to forest tree species yet. Therefore, we want to test Ĝ on European beech, a highly outcrossing species, to test if Ĝ provides insights into genotype-phenotype associations that would remain undiscovered with GWAS.

## Material and methods

### Genotypes and phenotypes

A data set of 514 individual trees was collected from five beech stands in the South-Eastern Romanian Carpathians along an altitudinal gradient associated with changes in precipitation and temperature (Grigoriadou-Zormpa et al. 2024). The exact geographical locations and more details about the stands are reported by Grigoriadou-Zormpa et al. (2024). The beech stands are referred to as Lempes, Tampa, Solomon, Lupului and Ruia. In Ruia, 110, in Lupului and Lempes 101, in Solomon 102 and in Tampa 99 trees were sampled. Lempes was the lowest elevation site at 550 to 600 m with the lowest mean annual precipitation of 712 mm, followed by Tampa located at 650-700 m with 791 mm, Solomon at 800 to 900 m with 855 mm and Lupului at 1000 to 1150 m with 957 mm mean annual precipitation. Ruia was the highest location at 1300 to 1450 m with the highest mean annual precipitation of 1023 mm. The tree stands are naturally regenerating and were established more than 100 years ago (Grigoriadou-Zormpa et al. 2024).

The phenotypic variables comprised four different traits, which were leaf nitrogen and leaf carbon content, carbon isotope composition δ^13^C in leaf dry mass and stomatal density. The leaves of the sun-exposed crowns of the trees were collected in August 2020, 2021 and 2022. Stomatal density was measured by counting the stomata within a fixed area of 0.1938 mm^2^ (Yücedağ et al. 2019) on the abaxial side of one dried sun exposed leaf per tree sampled in 2020. The stomata were counted based on a stomatal imprint created with fingernail polish and tape from dried leaves (Yücedağ et al. 2019). Carbon isotope composition (δ^13^C) in leaf dry mass, leaf nitrogen and carbon content were measured in 2020, 2021 and 2022. To evaluate changes in carbon isotope composition (δ^13^C) in leaf dry mass, leaf nitrogen and carbon content, homogenized plant material was analyzed by the Centre for Stable Isotope Research and Analysis at the University of Goettingen by elemental analysis combined with isotope ratio mass spectrometer measurements (Werner et al. 1999). Additionally, we calculated the C/N ratio based on the molar content [mmol/g] and molar weights of carbon and nitrogen. Stomatal closure due to reduced soil water availability and altered plant water status reduces the concentration of CO_2_ in the stomatal cavities and the concentration of the heavier ^13^C isotope increases compared to ^12^C (Fotelli et al. 2003). Consequently, plant water potential can be related to carbon isotope composition (δ^13^C) (Fotelli et al. 2003). δ^13^C is additionally influenced by changing CO_2_ levels and photosynthetically active radiation, so this trait shows strong variability across the growing season (Fotelli et al. 2003).

We removed extreme phenotypic outliers. The outliers and distribution of trait measurements are shown in supplemental file S1. Additionally, we examined the correlations of the traits with each other, which are shown in supplemental file S2. The correlations of the traits measured in different years are shown in supplemental file S3. Correlations were analyzed using R version 4.3.1 (R Core Team 2024).

Individual trees were sequenced by using the single primer enrichment technology (SPET) (Scaglione et al. 2019). This sequencing method targets SNPs located within and close to genes and flanking regions of ±2 kb (Scaglione et al. 2019). The target regions were determined by previous whole-genome sequencing of a subset of 96 individual trees by IGATech (IGA Technology Services Srl, Udine, Italy). Reads were aligned to the *F. sylvatica* reference genome version 2 (Mishra et al. 2022) by using BWA-MEM v0.7.17 (Li and Durbin 2009). Only aligned reads with a mapping quality ≥10 were kept and duplicated reads were removed. Variant calling was done along with filtering for minimum read number of an allele in a sample with Freebayes v1.3.6 (Garrison and Marth 2012; Grigoriadou-Zormpa et al. 2024). Normalization and filtering for read depth > 6 and < 257 were performed using bcftools (Danecek et al. 2011; Grigoriadou-Zormpa et al. 2024). After this, a total of 838,522 SNP markers were kept. The data set was filtered by PLINK for minor allele frequency (MAF) > 0.194% (which results in at least two observations at a marker), missingness of ≤ 10% at each SNP marker and for SNP markers without chromosome annotations. The final data set contained 758,465 SNP markers and 514 individuals used for GWAS. Not for all genotyped individual trees phenotypic measurements were available. For stomatal density, 314 individual trees were phenotyped. Four hundred forty individual trees were phenotyped for the δ^13^C measurements in 2020, 457 in 2021 and 454 in 2022. For the leaf nitrogen content measurements, 451 individual trees were phenotyped in 2020, 457 in 2021 and 450 in 2022. For the C/N ratio, 440 individual trees were phenotyped in 2020, 454 in 2021 and 450 in 2022.

### GWAS

To test for associations between SNPs and traits, we used a linear model in PLINK v.1.90 (Purcell et al. 2007). All 10 phenotypic variables were tested separately. The first two main components of the principal component analysis (PCA) based on the neutral markers were included as covariates to account for population structure. Neutral SNP markers were identified by Grigoriadou-Zormpa et al. (2024) for population genetic analysis. To identify the neutral SNP markers Grigoriadou-Zormpa et al. (2024) determined intergenic regions by using SnpEff (version 4.3t, Cingolani et al. 2012). Afterwards the filtering based on LD and missingness <10% was done, which resulted in a data set of 154,437 putatively neutral SNPs. To account for environmental effects during sampling in 2020, 2021 and 2022, the first principal coordinate based on precipitation and temperature during the growing season from April until August (supplemental Fig. S4 and supplemental Table S5) in the year of sampling was included (supplemental Fig. S6) in the GWAS for δ^13^C. This was done because δ^13^C shows strong variability across the growing season (Fotelli et al. 2003). Temperature and precipitation weather data were downloaded from Copernicus Climate Data Store (CDS 2024) for 2020, 2021 and 2022 (available in supplemental Fig. S4, Table S5 and Fig. S6). The principal coordinate analysis (PCoA) was performed on monthly temperature and precipitation measured from April until August for the years separately using the *cmdscale()* R function in the R version 4.3.1 (R Core Team 2025).

The significance thresholds were derived based on a permutation test using the same set of SNP markers (Purcell et al. 2007). The permutation test was conducted in the adaptive permutation mode, which can terminate before completing all possible iterations, when there are no changes anymore (Purcell et al. 2007). This adaptive permutation mode was run with ten minimal and 1,000,000 maximal permutations. The level at which no markers were identified as significant in our permutation test, was declared the significance threshold, which in our case was p ≤ 0.000001, which was the same for all traits. QQ plots are shown in supplemental file S7.

### Gene variant annotation

SnpEff (version 4.3t, Cingolani et al. 2012) was used for gene variant annotations. An annotation data base was built based on the *F. sylvatica* reference genome (Mishra et al. 2022), which was then also used in this study for the gene variant annotation. SnpEff uses this data base and the VCF file from this study to annotate gene variants, their effects and predicted impacts (Cingolani et al. 2012). The effects comprise missense variants, variants causing stop codons, frameshift variants, splice region variants and several more (Cingolani et al. 2012). A missense variant causes a codon change that produces a different amino acid (Cingolani et al. 2012).

### Ghat

Groups of stands growing under different environmental conditions were treated as separate populations to implement Ĝ (Fig. 1). The groups were formed on the basis of altitude and the amount of precipitation (Fig. 1). Group 1 included the stands Ruia and Solomon which were located at altitudes > 800 m and with a mean annual precipitation > 850 mm (Fig. 1). Group 2 included the stands Tampa and Lempes, which were all located at altitudes < 800 m and with a mean annual precipitation < 800 mm (Fig. 1). Lupului was removed from the Ĝ analysis, because of the age of the stand in comparison to the other stands (Fig. 1). The marker effect estimates were calculated, by using the rrBLUP R package (Endelman 2011).

**Fig. 1:**
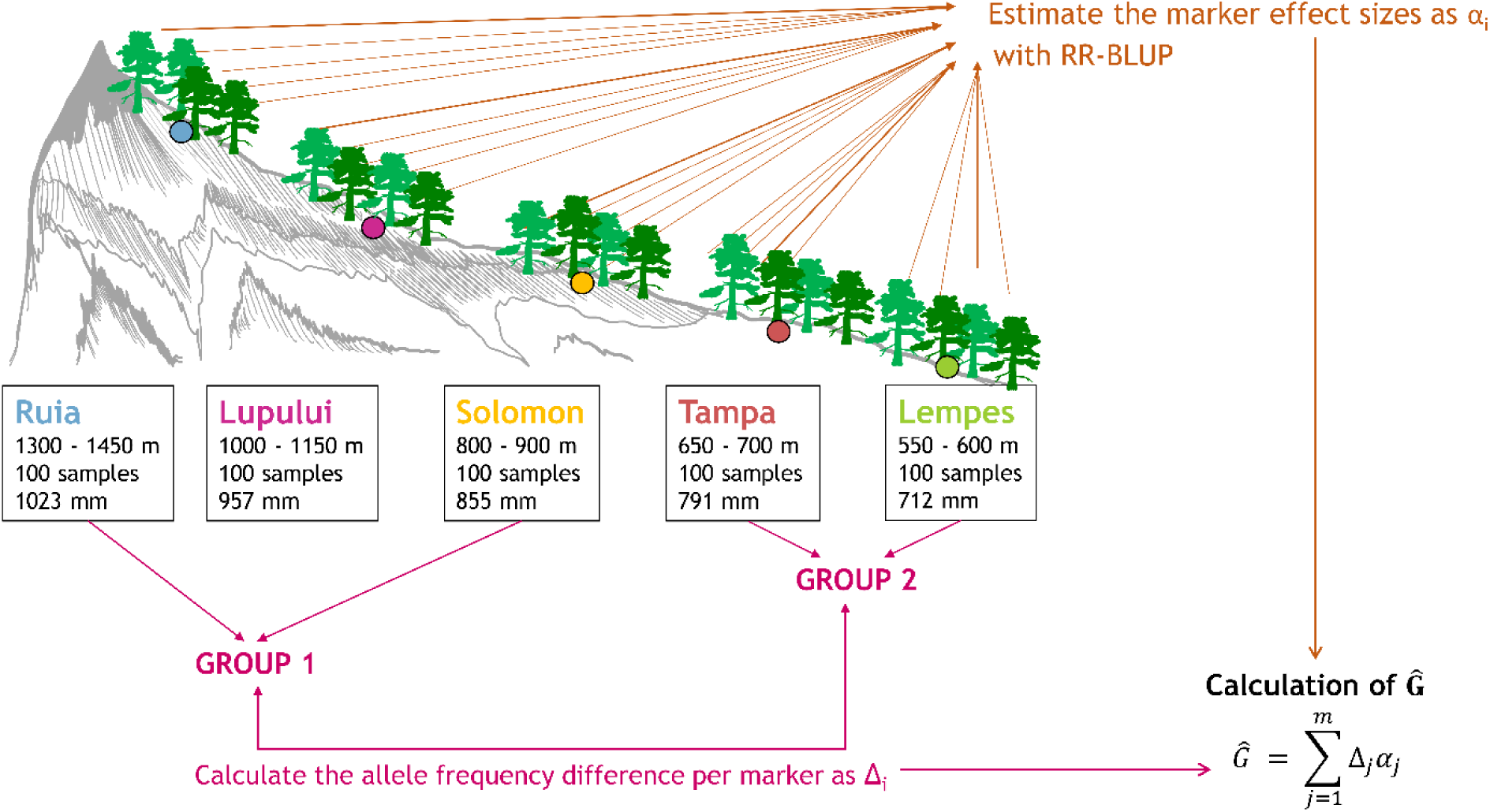
Simplified scheme of the location of the beech stands along the altitudinal gradient and the calculation of Ĝ by evaluating the marker effect sizes and allele frequency differences between groups of different environmental conditions across the entire genome.

Filtering for minor allele frequency (MAF) and missingness <0.05 was done before missing observations were imputed based on the mean allele frequency of the stands observed at each marker in R version 4.3.1 (R Core Team 2024). When Ĝ was calculated for δ^13^C, the weather data of the different years of sampling (see supplemental Figure S4 and Table S5) was included in the model to estimate the marker effects. Ĝ was calculated with the Ghat R package (Mahmoud et al. 2023). The number of permutations was set to 1000. The number of effective markers was first calculated based on a function called “ld_decay”, which is provided by the Ghat package (Mahmoud et al. 2023). To evaluate its impact on Ĝ, we additionally tested 100 randomly generated traits. These 100 random traits were generated by simulating a normal distribution with a mean of 25.93 and a standard deviation of 3.67 (mean and standard deviation of the trait leaf nitrogen content), then the traits were randomly associated to the individual trees. In the test runs we also tested if these random traits are identified as being under significant selection. When the random traits were identified as being under significant selection, they were considered to be false positive observations. Based on the 100 random traits, a rate of false positive observations was calculated and used to evaluate the number of effective markers. Later, in the actual Ĝ test, we tested the four different traits stomatal density, δ^13^C, leaf nitrogen content and C/N ratio to identify signals of polygenic selection.

### Cross-validation of marker effect sizes

To evaluate the marker effect estimates used in the Ĝ test, we conducted a cross-validation. For the cross-validation, the data set was randomly partitioned into five sets to be used in five-fold cross-validation with 100 replications (Endelmann 2011). In the cross-validation, the phenotypic values per trait of the testing set were predicted based on marker effect estimates which were calculated based on the training set (Endelmann 2011). The marker effect estimates were calculated with rrBLUP, using the same method as for the Ĝ test (Endelmann 2011). The predictive ability was calculated as correlation of predicted trait estimates and observed trait measurements for the different traits of the testing set (Endelmann 2011). Additionally, another cross-validation was conducted in which one stand was left out and the remaining stands were used for prediction. With this cross-validation we could assess how well the stands can predict a new stand.

### Calculation of linkage disequilibrium (LD) decay

We created a pairwise LD heat map with the R package LDheatmap for the regions which were associated with the traits and putatively under selection (Shin et al. 2006).

## Results

### GWAS revealed a peak of markers on chromosome 10 strongly associated with stomatal density and to a smaller extent with δ^13^C

Stomatal density was significantly associated with 121 markers, of which 101 are located on chromosome 10 (Fig. 2). The remaining markers are located on chromosomes 1, 4, 5, 6, 8 and 9, with one to five markers on each chromosome (Fig. 2). The 101 markers on chromosome 10 are located between ∼4.89 to ∼13.67 Mb, which appear as one wide peak (Fig. 2).

**Fig. 2:**
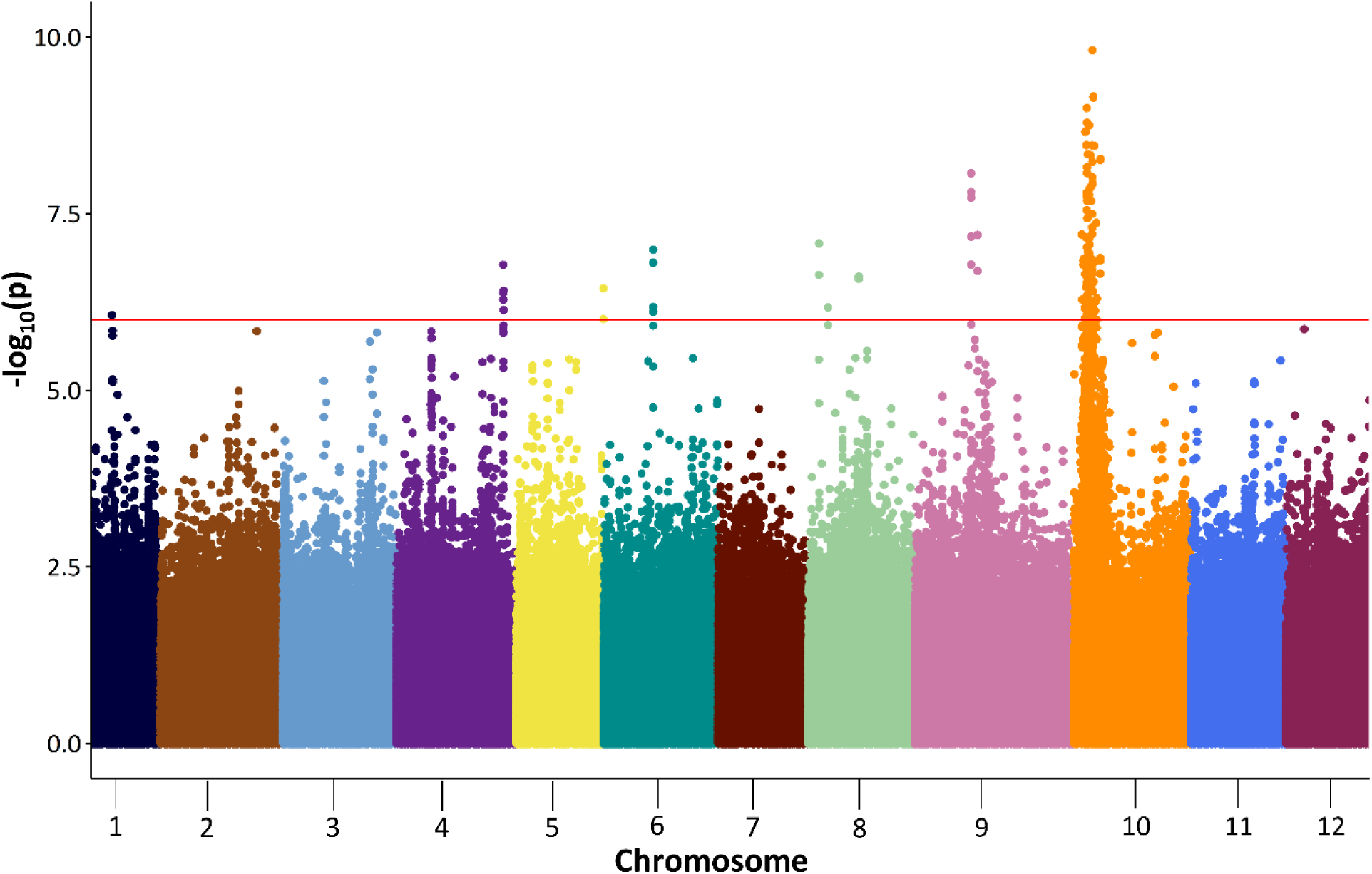
Manhattan plot for stomatal density with premutation derived significance thresholds of p ≤0.000001 (red horizontal line) with p-values plotted as -log_10_(p) against the position in bp.

A similar peak on chromosome 10 appears in the GWAS for δ^13^C measured in 2020 and 2022 (Fig. 3). But this peak is smaller and contains almost no significant SNP markers (Fig. 4.3). Only for δ^13^C measured in 2020, two SNP markers exceeded the permutation derived significance levels of p ≤0.000001 (Fig. 3). No significant SNPs were associated with δ^13^C measured in 2021 and 2022 (Fig. 3).

**Fig. 3:**
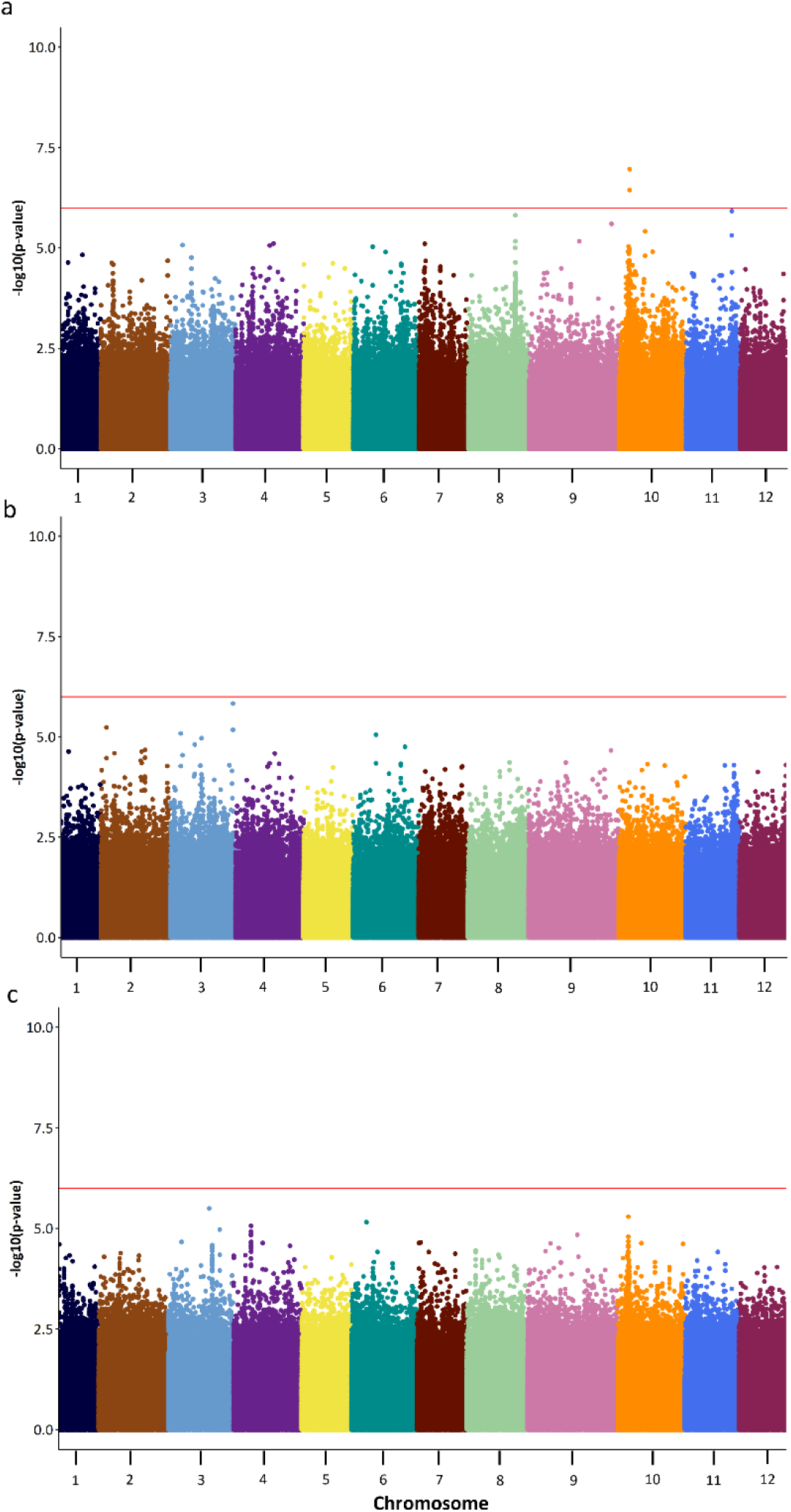
Manhattan plot for δ^13^C measured in 2020 (**a**), 2021 (**b**), and 2022 (**c**) with permutation derived significance thresholds of p ≤0.000001 (red horizontal line) with p-values plotted as -log10(p) against the position in bp.

**Fig. 4:**
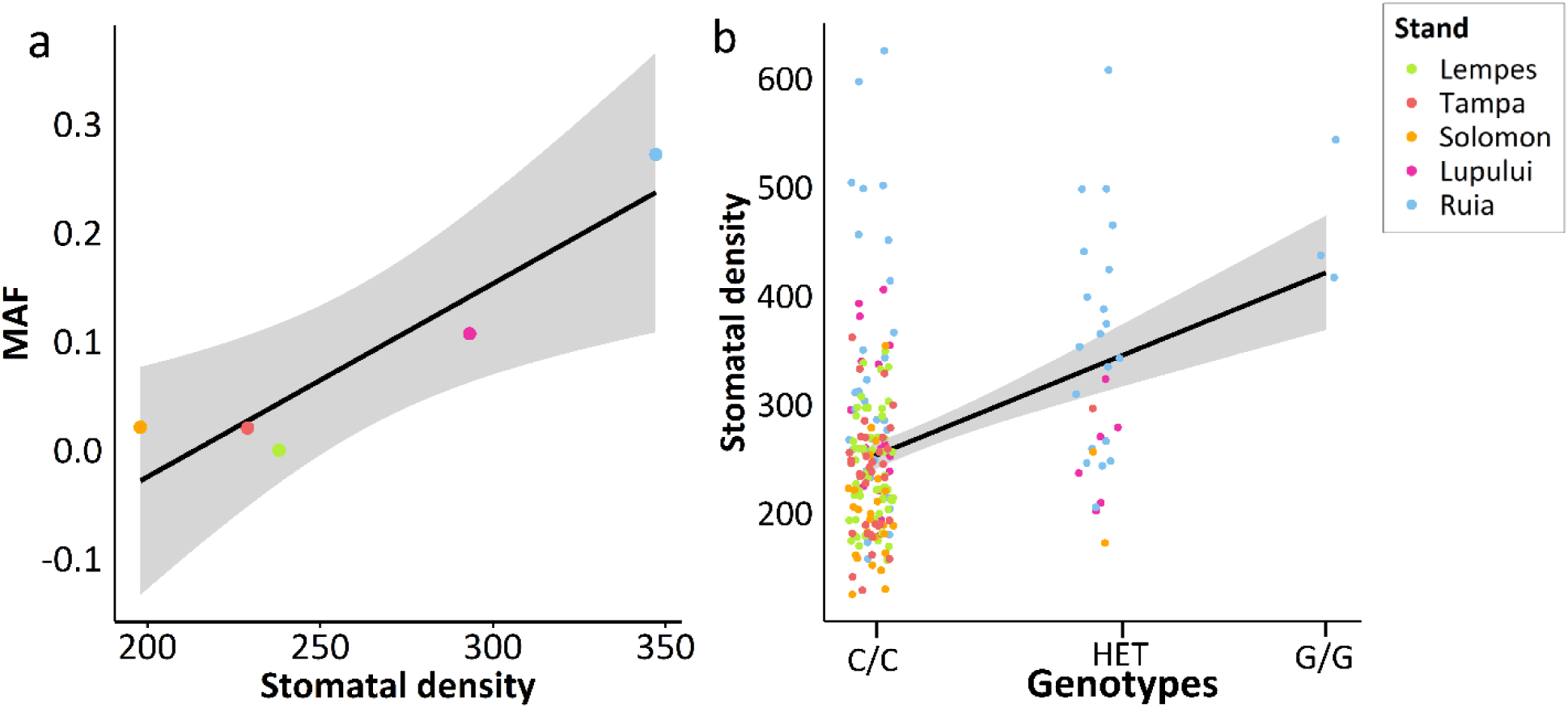
**a**) Correlation between stomatal density and the minor allele frequency (MAF) of the significant marker (p ≤0.000001) on chromosome 10 at ∼ 13.435 Mb with a correlation coefficient of 0.932 (p-value = 0.021) and **b**) Linear regression of marker genotypes against stomatal density with a coefficient of determination R^2^ of 0.1483 (p = 0.0001258). This marker was annotated with gene variant *Bhaga_10.g1651*, with the underlying gene *XP_023902123.1*, also described as *BONZAI1*.

Table 1 presents the most interesting GWAS results for stomatal density, focusing on candidate SNPs within 20 kb of gene annotations. The whole list of significant SNPs for stomatal density and δ^13^C measured in 2020 can be found in supplemental Table S8. The SNP markers significantly associated with δ^13^C measured in 2020 were not located within 20 kb of impactful parts of coding regions of genes (supplemental Table S8).

**Table 1:**
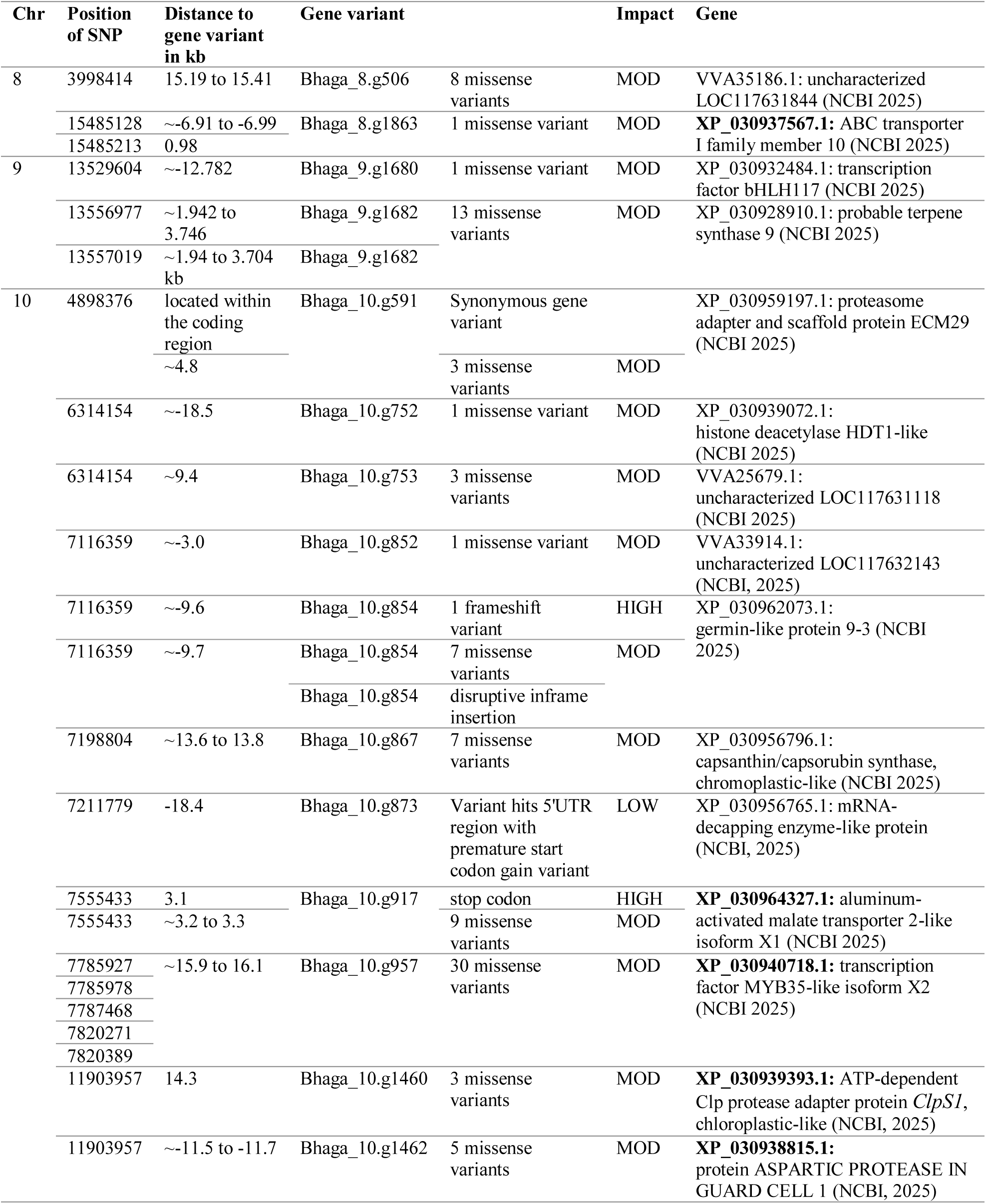
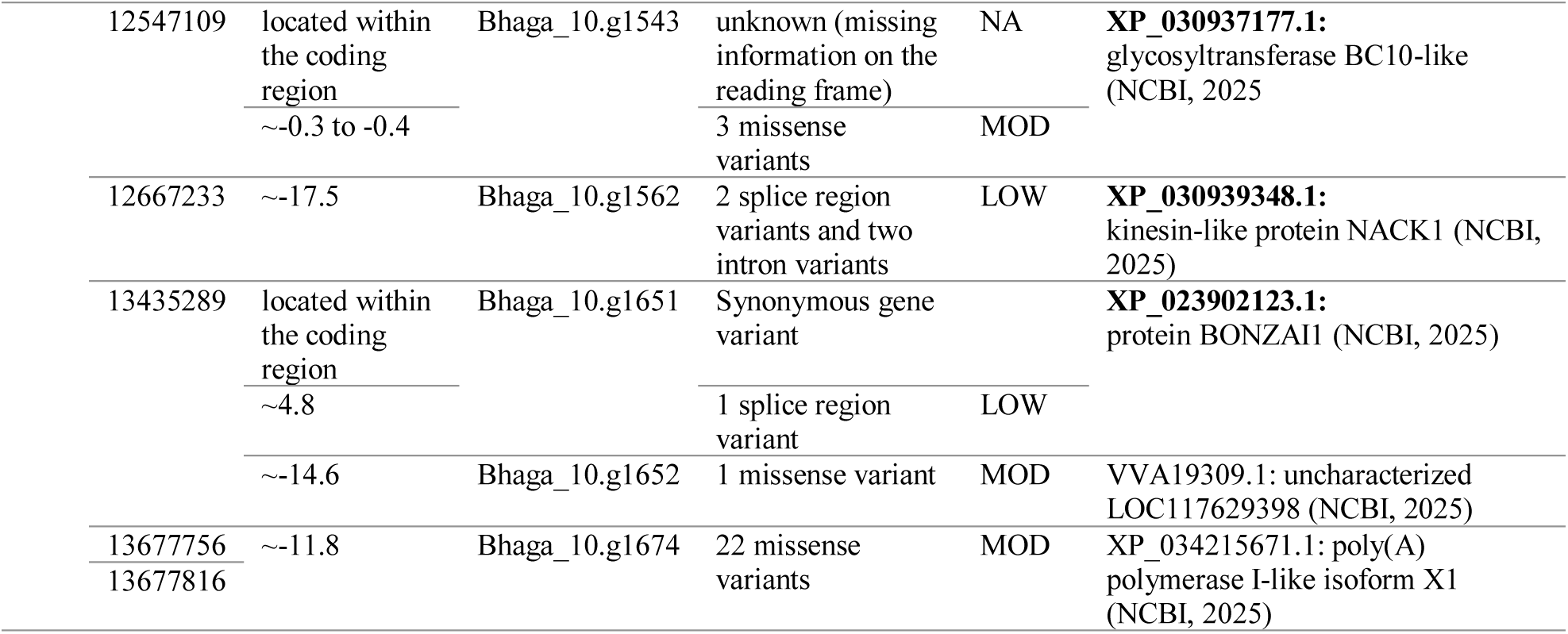
List of candidate SNPs with gene annotations associated with stomatal density with permutation derived significance levels of p ≤0.000001 and their annotated gene variants, the underlying genes with their locations, functions and predicted impact (HIGH, MOD (moderate) and LOW) based on SnpEff (Cingolani et al. 2012). Candidate genes involved in stomatal regulation, chloroplast biogenesis, and chloroplast metabolism are highlighted in bold face.

At the marker on chromosome 10 at ∼13.435 Mb, the MAF increases with increasing stomatal density which follows the altitudinal gradient associated with precipitation and temperature (Fig. 4). For all other markers on chromosome 10 from ∼4.89 to 13.67 Mb, we observed the same trend (see supplemental Table S9 and Fig. S10). In all cases, the minor allele frequency (MAF) is correlated with stomatal density (see supplemental Table S9 and Fig. S10).

### Associations of SNP markers with leaf nitrogen and carbon traits differed across years of sampling

Figure 5 displays the Manhattan plots for leaf nitrogen content and C/N ratio measured in 2020, 2021 and 2022, which were the other traits with significant associations. The SNP markers associated with the same trait measured in different years did not overlap (Fig. 5).

**Fig. 5:**
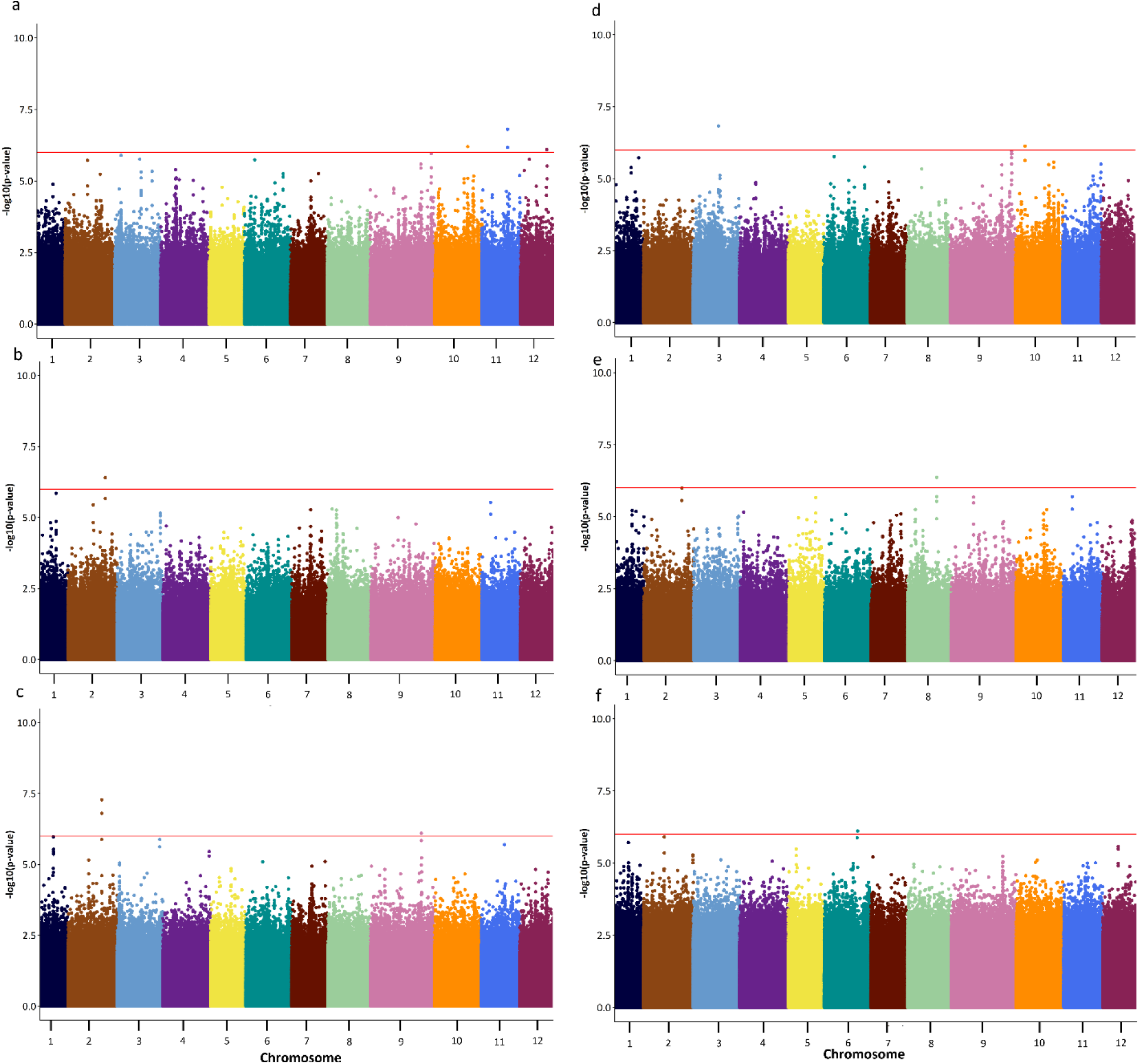
Manhattan plot for leaf nitrogen content measured in 2020 (**a**), 2021 (**b**), and 2022 (**c**) and C/N ratio measured in 2020 (**d**), 2021 (**e**), and 2022 (**f**) with permutation derived significance thresholds of p ≤0.000001 (red horizontal line) with p-values plotted as -log_10_(p) against the position in bp.

Table 2 shows the GWAS results for leaf nitrogen and carbon traits, which include only candidate SNPs of genes, which were < 20 kb away from the coding region of genes. These candidate SNPs were associated with leaf nitrogen content and C/N ratio measured in 2020, 2021, 2022 (Table 2).

**Table 2:**
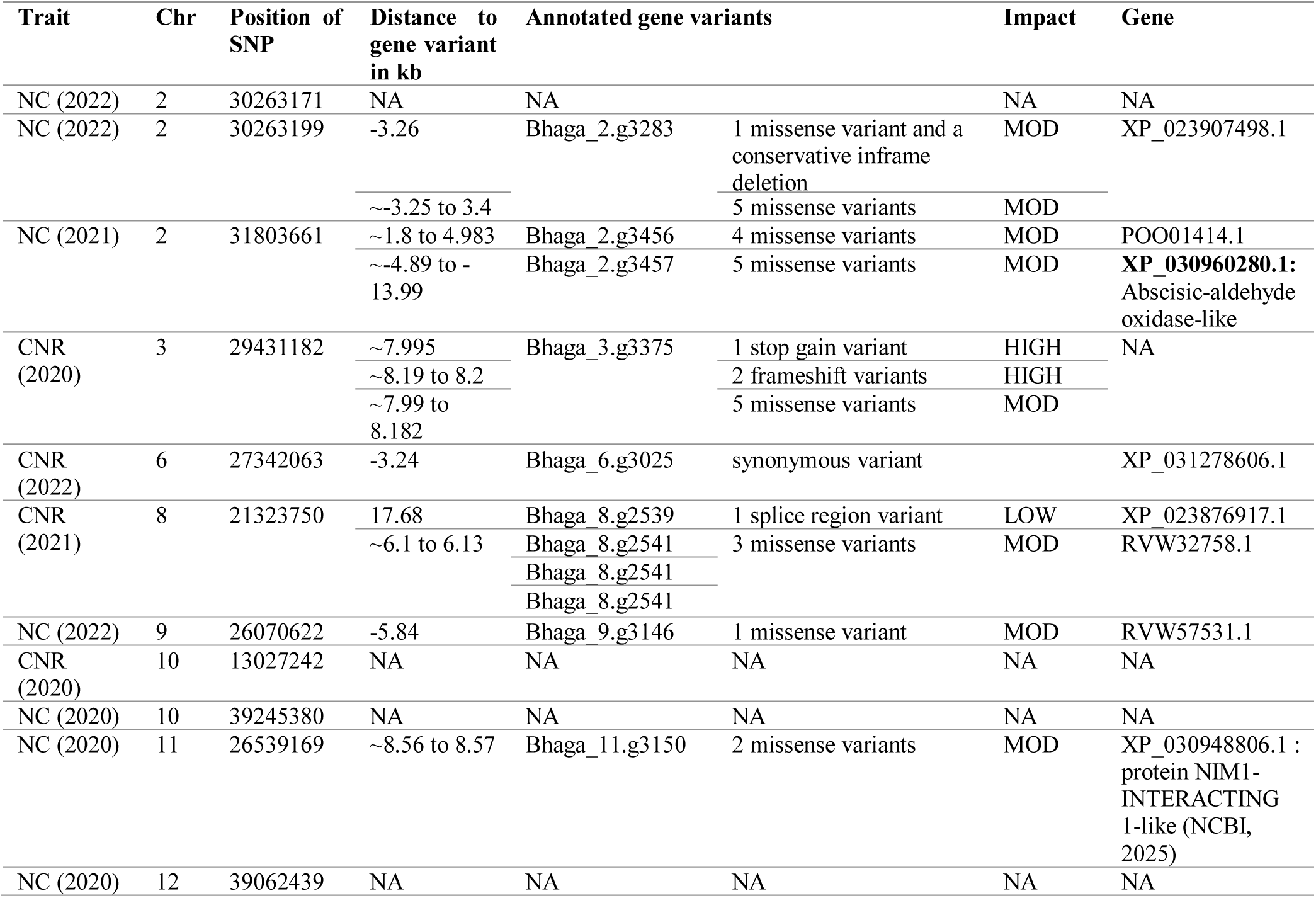
List of candidate SNPs associated with the other drought-stress related traits NC (leaf nitrogen content) and CNR (C/N ratio) with permutation derived significance levels of p ≤0.000001 and their annotated gene variants, the underlying genes with their locations, functions and predicted impact (HIGH, MOD (moderate) and LOW) based on SnpEff (Cingolani et al. 2012).

For C/N ratio and leaf nitrogen content, we do not observe a consistent trend for the significant markers (see supplemental Table S10, Figure S12 and S13). Supplemental Figure S11 shows the correlations for MAF and leaf nitrogen content in 2021 at the marker on chromosome 2 at ∼31.804 Mb, which was annotated with Bhaga_2.g3457 (*abscisic-aldehyde oxidase-like)*.

### Ghat indicates signals of positive polygenic selection for leaf δ^13^C

To evaluate the calibration of Ĝ, we tested 100 randomly generated traits. The calculated number of effective markers estimated based on LD decay over the entire genome was 838,522, which corresponds to the number of total markers. This resulted in 90 out of our randomly generated traits appearing as under selection by the uncalibrated Ĝ test. We did a grid search for effective marker number (ranging from 1,000 to 800,000). With 7,000 effective markers, we observed a rate of false positive observations <10% (see supplemental file S14). Results from cross-validation of predictions calculated with marker effect estimates based on 100 replications are available in supplemental file S15.

We calculated Ĝ between the high precipitation (Ruia and Solomon) and low precipitation group (Tampa and Lempes) and observed positive selection with a correlation coefficient between allele effect estimates and allele frequency change of 0.1017 (p ∼0) and 0.0962 (p ∼0) for δ^13^C in 2020 and 2021 (see supplemental Table S16). Based on this observation, we can deduce that polygenic selection for δ^13^C decreased with increasing precipitation along the altitudinal gradient (see also supplemental Table S1). Ĝ calculated based on δ^13^C measured in 2022 was not significant (supplemental Table S16). Soil water availability and plant water status can be related to δ^13^C, because an increase in δ^13^C describes the reduced concentration of CO_2_ in the stomatal cavities under closed stomata to improve the plant water potential (Fotelli et al. 2003). Under these circumstances, water use efficiency is usually increased (Fotelli et al. 2003).

## Discussion

### Annotations of genes harboring SNP markers significantly associated with stomatal density underpin a functional role in stomatal patterning and regulation of leaf water status

Stomatal density was significantly associated with 121 different markers, of which 101 are located on chromosome 10. In total, 26 of these markers, are located between position ∼4.89 to ∼13.67 Mb on chromosome 10 and appear as one wide peak. Across this region we observed increased linkage disequilibrium in the stands Solomon, Tampa and Lupului between R^2^ of 0.5 to 0.6. A total of 16 genes was identified within this peak, five of which are reported to have a direct influence on stomata regulation, chloroplast metabolism, chloroplast biogenesis or plastid maintenance (Gou et al. 2015; Nishimura et al. 2015; Shen et al. 2018; Dabravolski and Isayenkov 2023; Wang et al. 2024; Frangedakis et al. 2024) in *Arabidopsis thaliana* L. Overall, we found gene annotations for 21 genes of candidate SNPs associated with stomatal density on four chromosomes, but most interesting genes were annotated for gene variants located on chromosome 10.

Another gene with a potential influence on stomata was annotated for two markers on chromosome 8 at ∼15.485 Mb. This gene is called *ABC transporter I family member 10* (NCBI 2024). The two markers on chromosome 8 at ∼15.485 Mb are 0.98 kb upstream from a missense variant in the coding region of this gene variant. *ABC transporter I family member 10* belongs to the family of ATP-binding cassette (ABC) transporter lipids (Voith van Voithenberg et al. 2019). In *Arabidopsis thaliana*, *ABC transporter I family member 10* was observed to be responsible for ABC protein generation (Voith van Voithenberg et al. 2019). Loss of this gene led to extremely dwarfed albino plants with impaired chloroplasts (Voith van Voithenberg et al. 2019). Strong LD was observed between the two markers on chromosome 8 at ∼15.485 Mb and the coding region of *Bhaga_8.g1863 (ABC transporter I family member 10)* (supplemental Fig. S17).

The genes, located on chromosome 10, with a direct influence on stomatal characteristics comprise *aluminum-activated malate transporter 2-like* (Dabravolski and Isayenkov 2023), *transcription factor MYB35-like* (Frangedakis et al. 2024)*, ATP-dependent Clp protease adapter protein ClpS1* (Nishimura et al. 2015)*, aspartic protease in guard cell 1* (Shen et al. 2018; Wang et al. 2024) and *BONZAI1* (Gou et al. 2015).

The gene variant *Bhaga_10.g917* corresponds to the underlying gene *aluminum-activated malate transporter 2-like (ALMT2)* (NCBI 2024). *Aluminum-activated malate transporter 2-like (ALMT2)* belongs to the aluminum-activated malate transporter (ALMT) gene family which is involved in tolerance to Al^3+^ and regulation of opening and closing of stomata in plants (Dabravolski and Isayenkov 2023). A stop codon of this gene is located ∼3.1 kb upstream from the marker found on chromosome 10 at ∼7.56 Mb that was associated with stomatal density (see supplemental Fig. S18). This stop codon occurs in the first third of the coding region of the gene variant (see supplemental Fig. S18). Furthermore, nine missense variants of the gene coding region are located ∼3.2 to 3.3 kb upstream from the marker. According to the pairwise LD (linkage disequilibrium) heat map analysis, we observed strong LD around the marker and this gene (see supplemental Fig. S18). All nine missense variants caused a codon change that resulted in a different amino acid sequence (see supplemental Table S19). At six missense variants a hydrophilic amino acid was replaced by a hydrophobic amino acid (see supplemental Table S19). The change from a hydrophilic amino acid to a hydrophobic amino acid can increase the stability of the sequenced proteins in vivo (Lindman et al. 2010). At two out of nine missense variants the amino acids phenylalanine and serine were replaced by leucine (see supplemental Table S19).

The gene variant *Bhaga_10.g957* corresponds to the underlying gene *transcription factor MYB35-like* (NCBI 2024). MYB related transcription factors like *transcription factor MYB35-like* control chloroplast biogenesis and target genes for carbon fixation and light harvesting (Frangedakis et al. 2024). Mutants of MYB-related genes appear bleached and contain small chloroplasts (Frangedakis et al. 2024). We found five markers on chromosome 10 at ∼7.78 to ∼7.82 Mb which were ∼16 kb upstream from 30 missense variants located within the coding region of *transcription factor MYB35-like* with a moderate impact on its expression (see also supplemental Fig. S20). Our pairwise LD heat map, available as supplemental Figure S20, shows that the significant marker is in very strong and almost complete LD (∼1) with the coding region of this gene. No change in the polar bonds was detected in 25 missense variants (see supplemental Table S19). At three missense variants a hydrophilic amino acid was replaced by a hydrophobic amino acid (see supplemental Table S19). Glutamic acid was replaced by lysin, alanine was replaced by valine, and arginine was replaced by glutamine (supplemental Table S19). In the remaining two missense variants, one hydrophobic amino acid was replaced by a hydrophilic amino acid (see supplemental Table S19).

For the gene variant *Bhaga_10.g1460*, the annotated gene *ATP-dependent Clp protease adapter protein ClpS1* also referred to as *ClpS1* was found (NCBI 2024). *ClpS1* forms together with *ClpF* an important unit in the chloroplasts which serves as an adaptor and is responsible for substrate recognition and delivery (Nishimura et al. 2015). Our identified marker on chromosome 10 at ∼ 11.9 Mb is located ∼14.3 kb downstream from three missense variants of the coding region of the gene variant *Bhaga_10.g1460* (*ClpS1)* (see also supplemental Fig. S21). At two missense variants of *Bhaga_10.g1460* (*ClpS1)*, a hydrophilic amino acid was replaced by a hydrophobic amino acid (see supplemental Table S19).

The marker was located in the intergenic region between *Bhaga_10.g1460* (*ClpS1)* and *Bhaga_10.g1462* (*ASPG1*), but the pairwise LD is only increased in some parts along this region (see also supplemental Fig. S21). The gene variant *Bhaga_10.g1462* was annotated as the gene *aspartic protease in guard cell 1*, also known as *ASPG1* (NCBI 2024).

Stomata pores are surrounded by guard cells (Negi et al. 2013). *ASPG1* was found to be overexpressed under drought stress in *Arabidopsis* (Yao et al. 2012). The overexpression of the *ASPG1* gene increases the abscisic acid (ABA) sensitivity in stomatal guard cells and leads to reduced water loss in transgenic *Arabidopsis* plants (Yao et al. 2012). Shen et al. (2018) also observed that this gene plays a role in seed dormancy, seed longevity and seed germination in *Arabidopsis*. In general, aspartic proteases are involved in various biological processes like chloroplast metabolism, biotic and abiotic stress responses and reproductive development (Wang et al. 2024). Our identified marker on chromosome 10 at ∼11.9 Mb is also located ∼12 kb downstream from five missense variants of *Bhaga_10.g1462* (*ASPG1*) (see also supplemental Fig. S21). At three missense variants of *Bhaga_10.g1462* (*ASPG1*), a hydrophilic amino acid was replaced by a hydrophobic amino acid (see supplemental Table S19).

For the gene variant *Bhaga_10.g1651*, the gene *BONZAI1* was annotated (NCBI 2024). BONZAI genes control global osmotic stress responses, ABA accumulation and stomata regulation in plants (Chen et al. 2020). *BONZAI1* regulates Ca^2+^ signaling under osmotic stress in rice (Chen et al. 2020). In *Arabidopsis*, *BONZAI1* was found to also promote stomatal closure and ABA accumulation (Gou et al. 2015). ABA plays an important role in stomatal closure (Chen et al. 2022). The significant marker on chromosome 10 at ∼13.44 Mb is located within the coding region of *Bhaga_10.g1651* (*BONZAI1*) (synonymous gene variant) and ∼4.8 kb downstream from this marker is a splice region variant of this gene (see supplemental Fig. S22). The gene variant *Bhaga_10.g1652* is located ∼14 kb downstream from this marker, but no gene description was found for this gene variant (see supplemental Fig. S22). One missense variant was found in the coding region of *Bhaga_10.g1652*, which is ∼14.6 kb downstream of the significant marker on chromosome 10 at ∼13.44 Mb (supplemental Fig. S22).

The remaining genes are involved in disease resistance, degradation and regulation processes. A potential role in stomatal density could not be directly inferred from the literature and further studies are needed. These comprise *histone deacetylase HDT1-like*, *germin-like protein 9-3*, *capsanthin synthase chromoplastic-like*, *mRNA-decapping enzyme-like protein*, *glycosyltransferase BC10-like*, *kinesin-like protein NACK1* and *poly(A) polymerase I-like* (NCBI 2024). Their description is available in supplemental Table S24.

We assume that the genes *histone deacetylase HDT1-like* (*Bhaga_10.g752*) and *glycosyltransferase BC10-like* (*Bhaga_10.g1543*) could also be important for stomatal density, since they are involved in stem vascular development and cellulose biosynthesis. Chhetri et al. (2020) detected genes encoding glycosyltransferases in a GWAS for stomatal density. Our significant marker on chromosome 10 at ∼12.5471 Mb is located within the coding region of *Bhaga_10.g1543 (glycosyltransferase BC10-like)*, but at this position the impact on the coding of the gene is unknown (missing information on the reading frame) (see supplemental Fig. S23). We observed very strong LD (∼1) across the entire coding region of this gene (see supplemental Fig. S23).

In contrast, *Capsanthin synthase chromoplastic-like* does not seem to be important for stomatal density. The importance of the other genes for stomatal density is also difficult to deduce. It may be possible that some of these genes are hitchhiking, because strong joint selection is acting across the entire region on chromosome 10 from ∼4.89 to ∼13.67 Mb.

### Significant correlations of stomatal density with minor allele frequency and environmental parameters indicate a contribution to local adaptation

All markers located on chromosome 10 between ∼4.89 to ∼13.67 Mb show a positive correlation between stomatal density and MAF (Fig. 4, supplemental file S9 and S10). In this study, we observed increasing stomatal density with increasing altitude and precipitation and decreasing temperature. Hamanishi et al. (2012) observed in *Populus* that lower stomatal density restricts the number of sites for water loss, thereby reducing transpiration. Li et al. (2023) found that under drought stress, *Populus* trees exhibit lower stomatal densities but increased stomatal length and width and improved stomatal control mechanisms. In this study we observed a correlation of stomatal density with water use efficiency measured as δ^13^C measured in 2020 at 0.23 (p ≤ 0.005), in 2021 at 0.18 (p ≤ 0.005) and in 2022 at 0.18 (p ≤ 0.001) (supplemental Fig. S2). Overall, we found slightly more genes involved in stomatal control and regulation than in stomatal development. Stomatal density and stomatal conductance are usually positively correlated, in this study stomatal conductance was not measured (Franks et al. 2009).

We observed very similar correlations between stomatal density and MAF across all markers on chromosome 10 between ∼4.89 to ∼13.67 Mb (Fig. 4, supplemental file S9 and S10). This indicates joint selection for this entire region, in which at least five genes with a direct influence on stomata are located. The correlation of MAF and stomatal density is not increasing in full agreement with the position of the stands along the altitudinal gradient (Fig. 4, supplemental file S9 and S10). Solomon and Tampa have slightly lower stomatal density than Lempes. This might be explained by the steeper slopes in Solomon and Tampa than in Lempes, which may have led to less water being retained over the years in Solomon and Tampa in comparison to Lempes.

Figure 6 displays the observed haplotypes across the entire region on chromosome 10 between ∼4.89 to ∼13.67 Mb and the pairwise LD heat map within the different stands. Observed haplotypes across the entire region differ especially between Ruia and Lupului, Tampa and Lempes (Fig. 6A). Pairwise LD is increased in Lupului, Solomon and Tampa in comparison to Ruia (Fig. 6B).

**Fig. 6:**
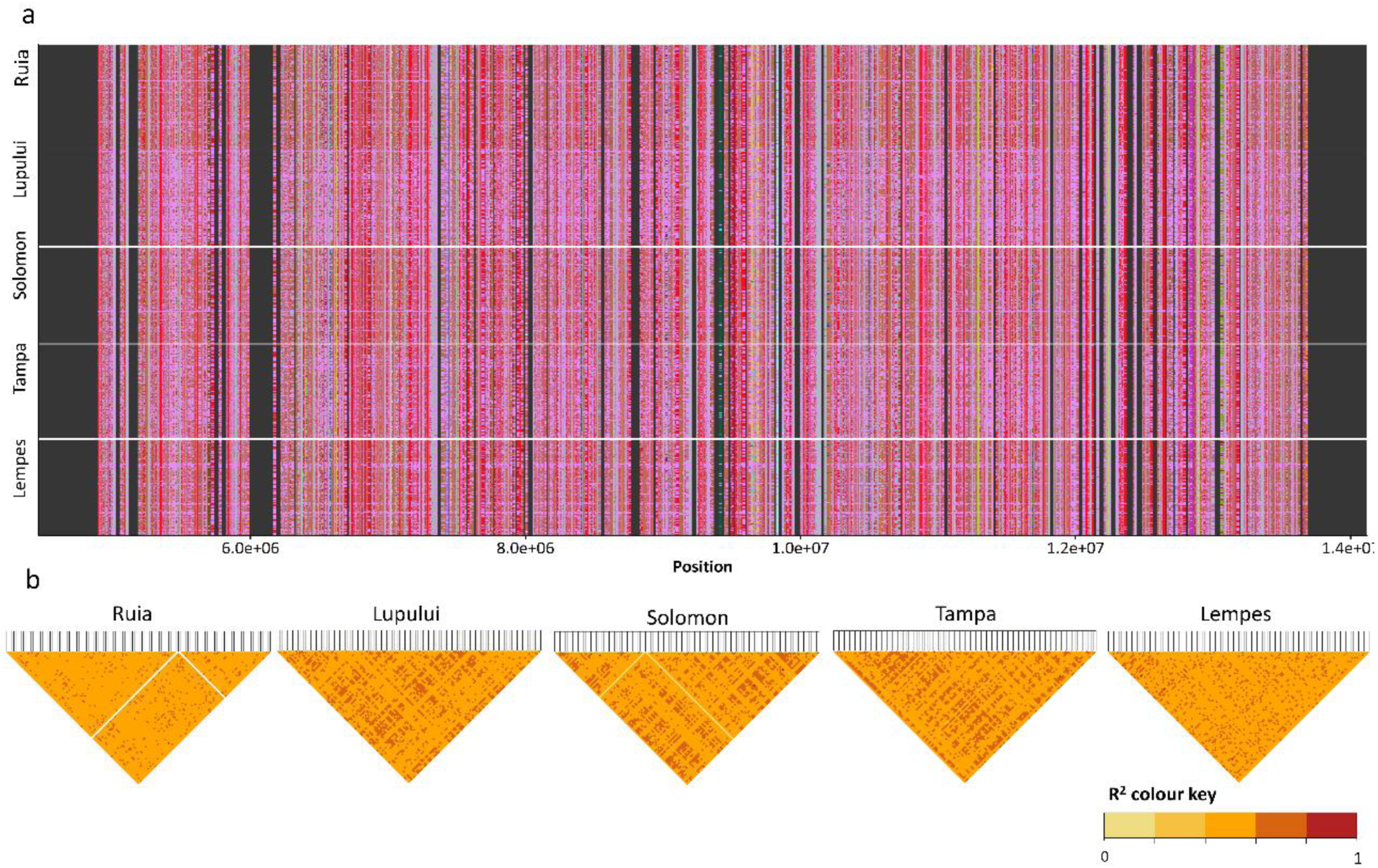
Region on chromosome 10 from ∼4.89 to ∼13.67 Mb and the observed haplotypes (**a**) and pairwise LD heat map for the different subpopulations (**b**). Further evidence supporting that this region on chromosome 10 is involved in local adaptation, is provided in Figure 7. *F_ST_*-based selection signature mapping revealed a peak on chromosome 10 from ∼4.512 to ∼47.742 Mb (Fig. 7). This peak is partly overlapping with the peak containing marker associations from the GWAS for stomatal density from ∼4.89 to ∼13.67 Mb on chromosome 10.

**Fig. 7:**
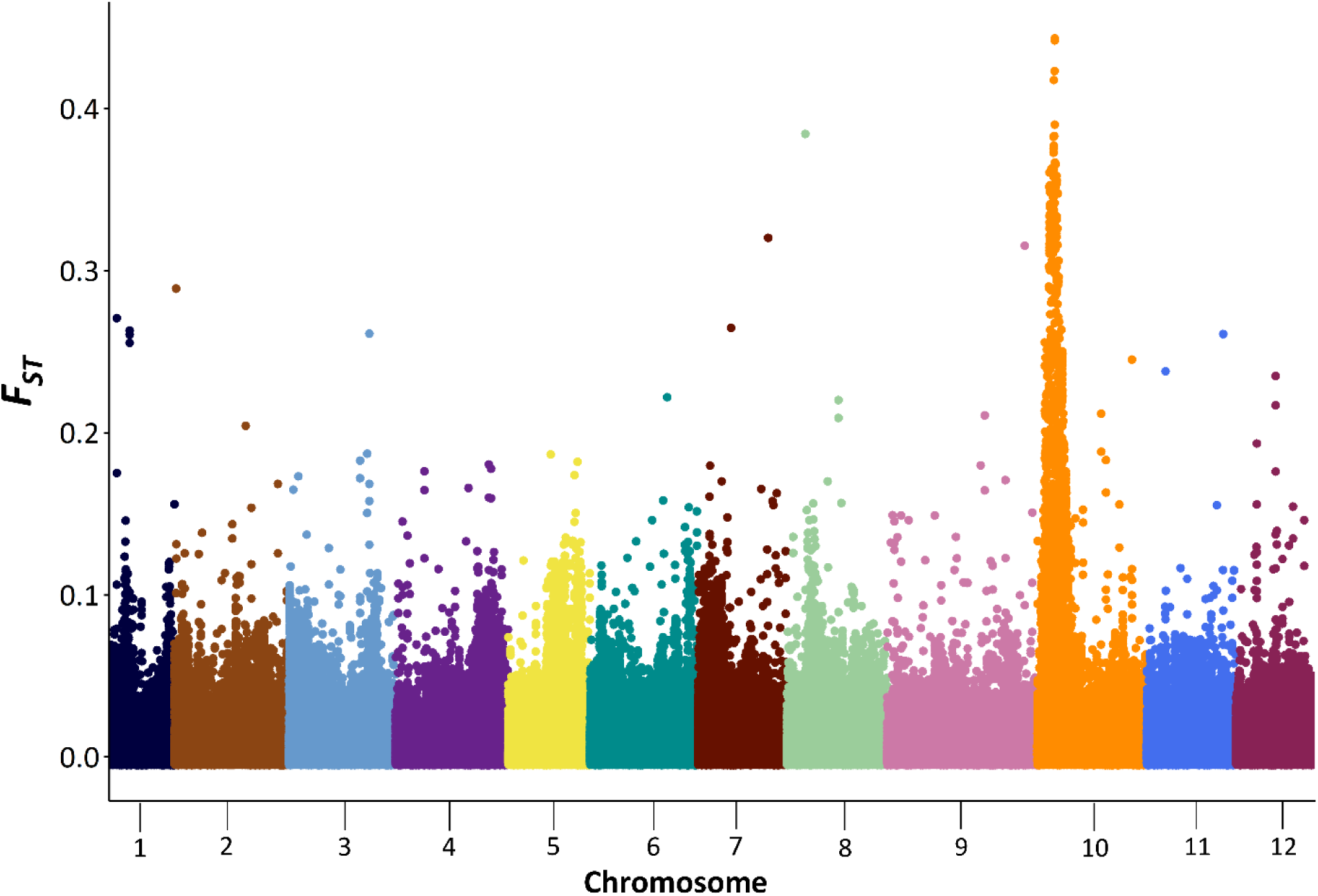
Genome-wide *F_ST_* values based in European beech (*Fagus sylvatica* L.) comprising 100 individuals adult beech trees per stand in five locations sampled in the South-Eastern Romanian Carpathians along an altitudinal gradient associated with precipitation and temperature.

Most GWAS for stomatal density identified several genes or QTL (Quantitative Trait Loci) (Bergmann et al. 2004; Gailing et al. 2008; McKown et al. 2019; Chhetri et al. 2020; Li et al. 2023). Li et al. (2023) also identified *Aluminum-activated malate transporter 12 (ALMT12)* in a GWAS and a co-expression network analysis in *Populus*. *ALMT12* belongs to aluminum-activated malate transporter (ALMT) family (Li et al. 2023). Based on our GWAS and the literature review, we can assume stomatal density has an oligogenic trait architecture and the identified region on chromosome 10 from ∼4.89 to ∼13.67 Mb seems to be very important in local adaptation. High peak SNP associations and an oligogenic trait architecture for stomatal density were also reported by McKown et al. (2019) and Chhetri et al. (2020) in *Populus trichocarpa* Hook. A linear regression of the different genotypes observed at the SNP on chromosome 10 at ∼7.82 Mb and stomatal density is available in supplemental Figure S25. The coefficient of linear regression calculated was considerably high (R^2^=0.2353, p ∼0).

All gene variants except for *Bhaga_10.g1674* were annotated based on other tree species of the family Fagaceae, which were *Quercus suber* L. and *Quercus lobata* Née (NCBI 2024). This makes the annotated gene variant results more reliable. Unfortunately, these genes were not studied in *Q. suber* L. and *Q. lobata* Née*. Bhaga_10.g1674* was annotated based on *Prunus dulcis* Batsch (NCBI 2024).

### GWAS results for leaf nitrogen and carbon varied across traits and year of sampling

In total, 11 markers were identified as being significantly associated with leaf nitrogen content and C/N ratio measured in 2020, 2021 and 2022. Eight of these markers were < 20 kb from coding regions of annotated genes (Table 2). In total, eight gene annotations for the ten markers were available (Table 2). We only found one underlying gene that was described in the literature. The gene variant *Bhaga_2.g3457* was annotated as *abscisic-aldehyde oxidase-like* (NCBI 2024), which was observed to be related to drought stress (Little and Eidt 1968). A marker on chromosome 2 at ∼ 31.8 Mb, near the abscisic aldehyde oxidase-like gene, was associated with leaf nitrogen content in 2021.

The gene *abscisic-aldehyde oxidase-like* is responsible for oxidation of abscisic aldehyde, which is the last step of abscisic acid (ABA) biosynthesis (Seo et al. 2000). ABA delays bud break in forest tree species and also inhibits simultaneously transpiration (Little and Eidt 1968). Peuke et al. (2002) observed an increase in ABA under stress in European beech. As mentioned before, ABA also plays an important role in stomatal closure (Chen et al. 2022). Lu et al. (2015) observed a connection between C/N ratio and ABA signaling pathways. Reduced leaf nitrogen supply can increase sensitivity of stomata (Gessler et al. 2017).

There is no consistent correlation trend between leaf nitrogen measured in 2021 and MAF (supplemental Fig. S11). MAF is increased in Lempes, Tampa (stands at low altitude) and Ruia (at high altitude), and decreased in Lupului and Solomon (supplemental Fig. S11). Leaf nitrogen content is decreased in Lempes, Tampa (stands at low altitude) and Ruia (at high altitude), and increased in Lupului and Solomon (supplemental Fig. S11). The phenotypic variability was observed to be much higher for traits like leaf nitrogen content and C/N ratio than for stomatal density (De la Torre et al. 2022). More stands along the altitudinal gradient and individuals per stand may have been useful to study polygenic traits with a high phenotypic plasticity like leaf nitrogen content and C/N ratio. The R^2^ value of the genotypes observed at the marker on chromosome 3 at ∼31.804 Mb and leaf nitrogen content measured in 2021 was 0.0069.

Other genes involved in ABA biosynthesis were *ASPG1* (*Bhaga_10.g1462*) and *BONZAI1* (*Bhaga_10.g1651*), which are close to or overlapping with SNP markers associated with stomatal density. Given the importance of ABA biosynthesis in drought tolerance, the marker on chromosome 2 at ∼31.8 Mb is a promising candidate SNP for drought stress tolerance.

No SNP marker was associated with more than one trait. However, associations for stomatal density and δ^13^C were observed in the same region on chromosome 10. We would have expected at least a few overlapping markers between leaf nitrogen content and C/N ratio, since C/N ratio is calculated based on the leaf nitrogen content. Also, no consistent association of markers with the same trait measured in different years was found. The correlation of C/N ratio measured in different years was relatively high with a correlation coefficient of 0.497 (p ≤ 0.0001), 0.460 (p ≤ 0.0001) and 0.510 (p ≤ 0.0001) (supplemental Fig. S2). The correlation of leaf nitrogen content measured in different years was similar (supplemental Fig. S2). Phenotypic variability of these traits may be too high to detect SNP-trait associations across years (De la Torre et al. 2022), which can also be observed in supplemental Figure S1. Additionally, leaf nitrogen content (N) and the C/N ratio are impacted by complex physiological mechanisms (Gessler et al. 2017). These complex physiological mechanisms depend on the C and N availability before the drought event (Gessler et al. 2017). Lower nutrient availability at the beginning of drought stress leads to a higher C/N ratio during drought, while higher nutrient availability leads to a lower C/N ratio during drought (Gessler et al. 2017). Short and intense drought stress usually causes higher nitrogen depletion, while long drought stress causes C starvation (Gessler et al. 2017). These complex physiological mechanisms may make association studies for these traits even more complicated.

Two SNP markers were associated with δ^13^C measured in 2020 at the permutation derived significance levels of p ≤0.000001. Only two SNP markers were identified as being associated with δ^13^C measured in 2021 and 2022. In the GWAS for δ^13^C measured in 2020 and 2022, we observed a peak on chromosome 10 that is overlapping with the peak observed in the GWAS for stomatal density on chromosome 10 from ∼4.89 to ∼13.67 Mb (Fig. 1 and Fig. 2). This observation is meaningful as δ^13^C captures changes in carbon isotope composition due to stomatal closure due to reduced soil water availability and plant water status (Fotelli et al. 2003).

### Ĝ results for δ ^13^C

The fact that only in one year two SNPs were associated with δ^13^C does not mean, there is no genetic basis for this trait. Previous studies showed, that it is difficult to identify significant markers for highly polygenic traits with infinitesimal sites contributing to the genetic variation (Beissinger et al. 2018; Zeng et al. 2018; Morales et al. 2021; Mahmoud et al. 2023). Therefore, we additionally applied the polygenic test Ĝ to our data set. According to Ĝ, δ^13^C in 2020 and 2021, was under highly significant polygenic selection (p ∼0) (see supplemental Table S16). δ^13^C decreased with increasing precipitation along the altitudinal gradient (p ∼0) in the two years (2020; 2021). Ĝ tested for δ^13^C measured in 2022 was not significant (see supplemental Table S16). The Ĝ results for 2020 and 2021 are in concordance with the literature. δ^13^C, which can be used as an indicator of plant water potential, is increased (or less negative) under heat and drought stress. This leads to reduced water loss under heat stress, which was also observed in previous studies (Querejeta et al. 2003; Fotelli et al. 2003).

In the GWAS for δ^13^C measured in 2020 and 2022, a peak on chromosome 10 which overlapped with the peak observed in the GWAS for stomatal density chromosome 10 from ∼4.89 to ∼13.67 Mb (Fig. 1 and Fig. 2). To summarize, we observed the clearest results for δ^13^C measured in 2020, with signals of positive polygenic selection and two significant SNP markers near multiple candidate SNPs for stomatal density. The MAF observed at these SNP markers on chromosome 10 at ∼ 10.9071 Mb and at ∼10.9073 Mb correlated positively with δ^13^C measured in 2020 at *|r|* ∼ 0.4910 (p-value = 0.08) and *|r|* ∼ 0.4843 (p-value = 0.06), but this correlation was not significant.

## Conclusions

Our research gives insights into the genetic architecture of different drought stress traits. We identified several very interesting candidate SNPs for stomatal density. Most of these candidate SNPs are located in one wide region on chromosome 10 from ∼4.89 to ∼13.67 Mb. Within this region, we found based on a literature review, annotations for five genes with a direct impact on stomatal density, which comprise *aluminum-activated malate transporter 2-like*, *transcription factor MYB35-like, ATP-dependent Clp protease adapter protein ClpS1, Aspartic protease in guard cell 1* and *BONZAI1.* The minor allele frequencies observed at the candidate SNP markers in this region are positively correlated with stomatal density, altitude and precipitation and negatively with temperature. Therefore, we can deduce that this entire region is important for local adaptation. Furthermore, we identified one interesting candidate SNP associated with leaf nitrogen content. For δ^13^C measured in one year, we identified two significant SNP marker associations, also located within the wide peak region on chromosome 10 from ∼4.89 to ∼13.67 Mb associated with stomatal density. We also observed signals of highly polygenic selection for δ^13^C. In conclusion, our research demonstrates that polygenic tests like Ĝ and GWAS can complement each other. Ĝ focuses on the small effect loci and uncovers their combined effect on a trait. Our GWAS worked very well regarding the identification of candidate SNP markers for stomatal density. We also identified a few SNP markers associated with leaf nitrogen content and C/N ratio. Unfortunately, there was no overlap between markers associated with the same trait measured in different years. Our GWAS did return only a limited number of candidate SNP markers associated with δ^13^C in one year, but the polygenic test Ĝ returned signals of polygenic selection for this trait. Our results suggest that traits like δ^13^C are polygenic, whereas stomatal density is influenced by a few larger-effect loci. Therefore, genomic prediction seems to be a promising tool for highly polygenic traits like δ^13^C. When traits like stomatal density are under selection, marker-assisted selection (MAS) of our few candidate SNPs for stomatal density may be sufficient. But for future genomic prediction or Ĝ studies, we would recommend the usage of larger data sets in terms of individuals, because we observed low predictive abilities in our cross validations based on this data set.

## Availability of data and materials

All the data and statistics about the present study have been included in the current manuscript in the form of figures and tables or in the supplemental material. Raw data is publicly available at figshare: 10.6084/m9.figshare.28748924. Our analysis code is publicly available on github: https://github.com/MilaTost/DroughtMarkers.

## Acknowledgements

We would like to thank all people involved in this project. We are thanking Jonah Fels, Jürgen Scheuring, Nikolas Heeger, Luca Schwendemann, Merle Kleikamp, Timo Unbehau, Jonas Daniel, Katharina Ziesing, Côme Sylvain-dutillow, Tania Dominguez-Flores, Christoph Leuschner, Katharina B. Budde, Mehdi Ben Targem, Thi Mai Trang Schneider, Mihnea Ciocirlan and Elena Ciocirlan.

