## Supplemental File for "Genome-wide association study in European beech (*Fagus sylvatica* L.) for drought stress traits"

**Supplementary files**

**S1:** Phenotypic trait measurements: diameter at breast height (DBH), stomatal density, leaf carbon content measured in 2020, 2021 2022, leaf nitrogen content measured in 2020, 2021 2022, C/N ratio calculated for 2020, 2021 2022 and water use efficiency as $\delta^{13}$C measured in 2020, 2021 2022.

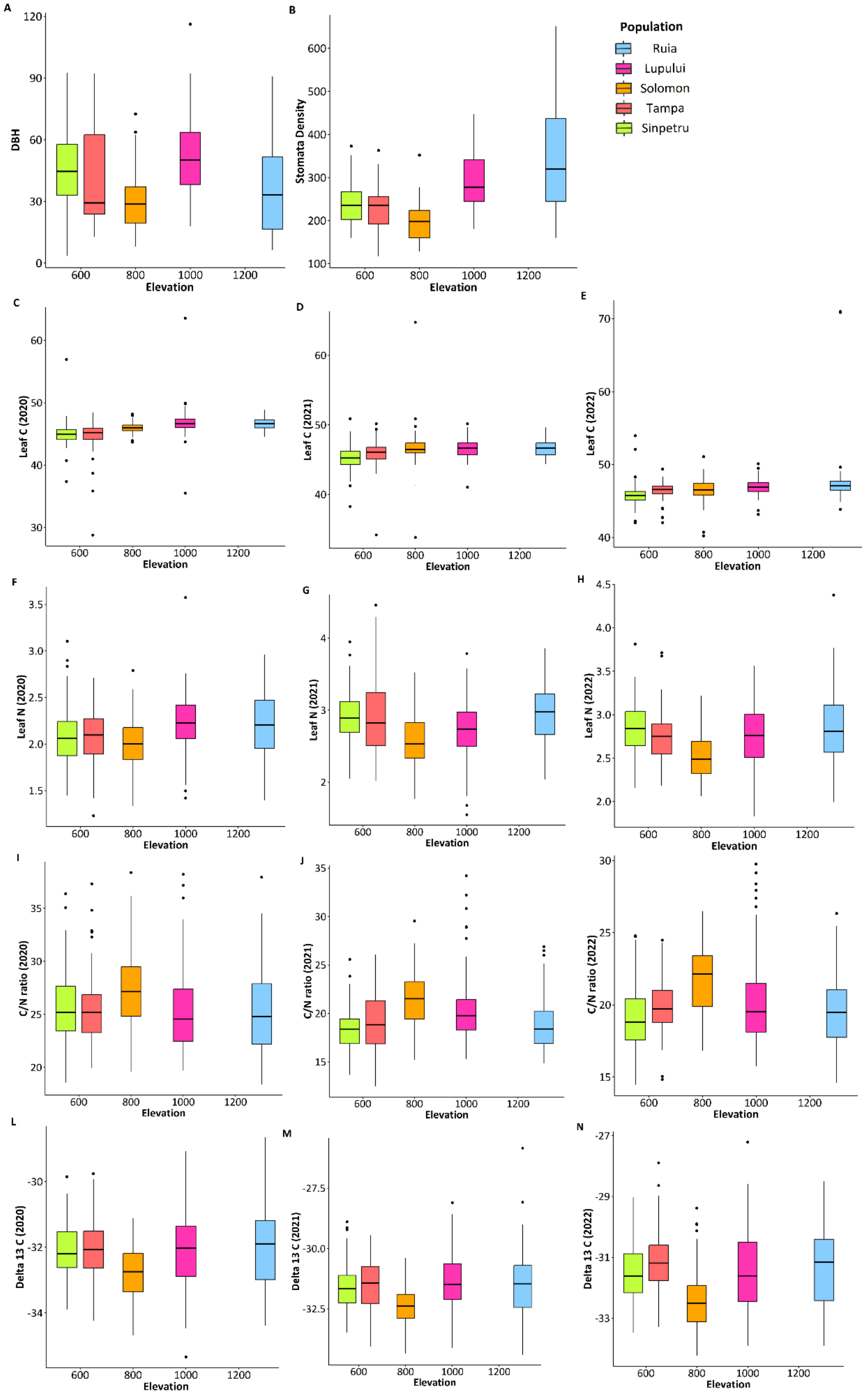

**S2:** Correlations of all phenotypic traits and their correlation coefficients (p ≤ 0.05).

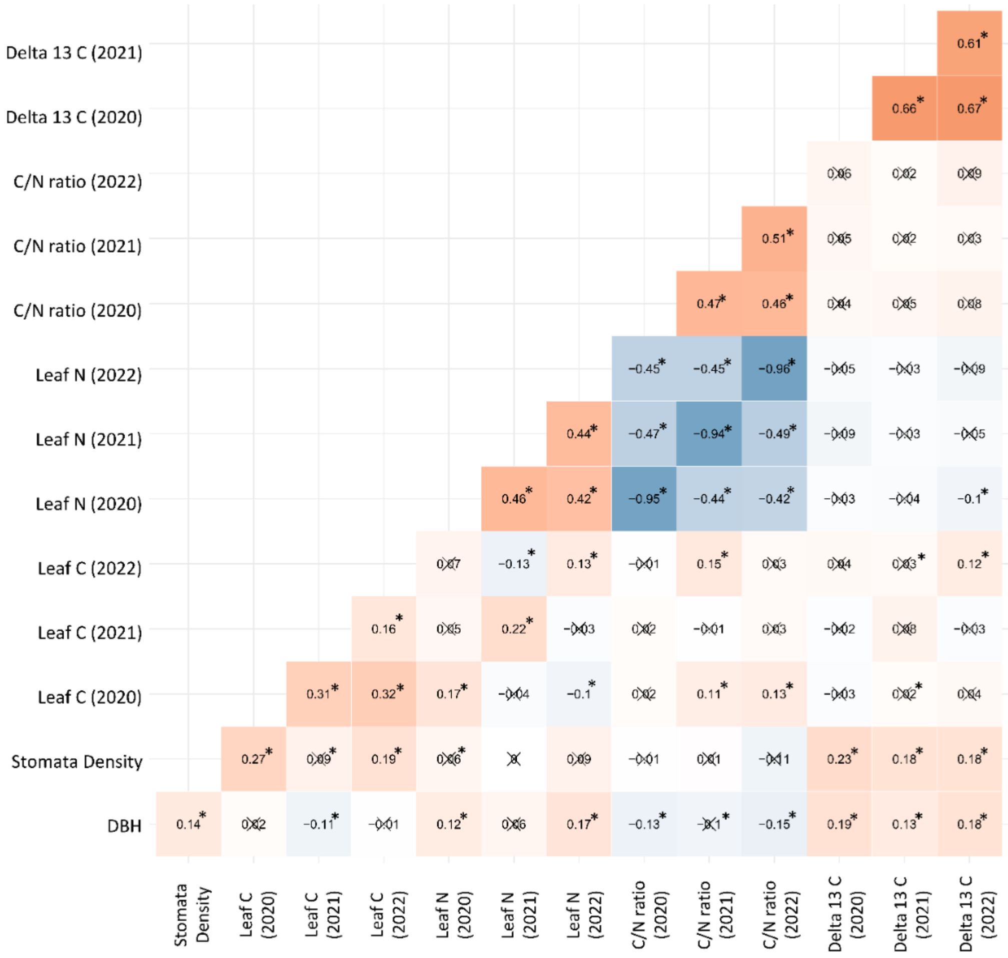

**S3:** Correlations of the traits measured in different years for leaf carbon content measured in 2020, 2021 2022, leaf nitrogen content measured in 2020, 2021 2022, C/N ratio calculated for 2020, 2021 2022 and water use efficiency as $\delta^{13}C$ measured in 2020, 2021 2022.

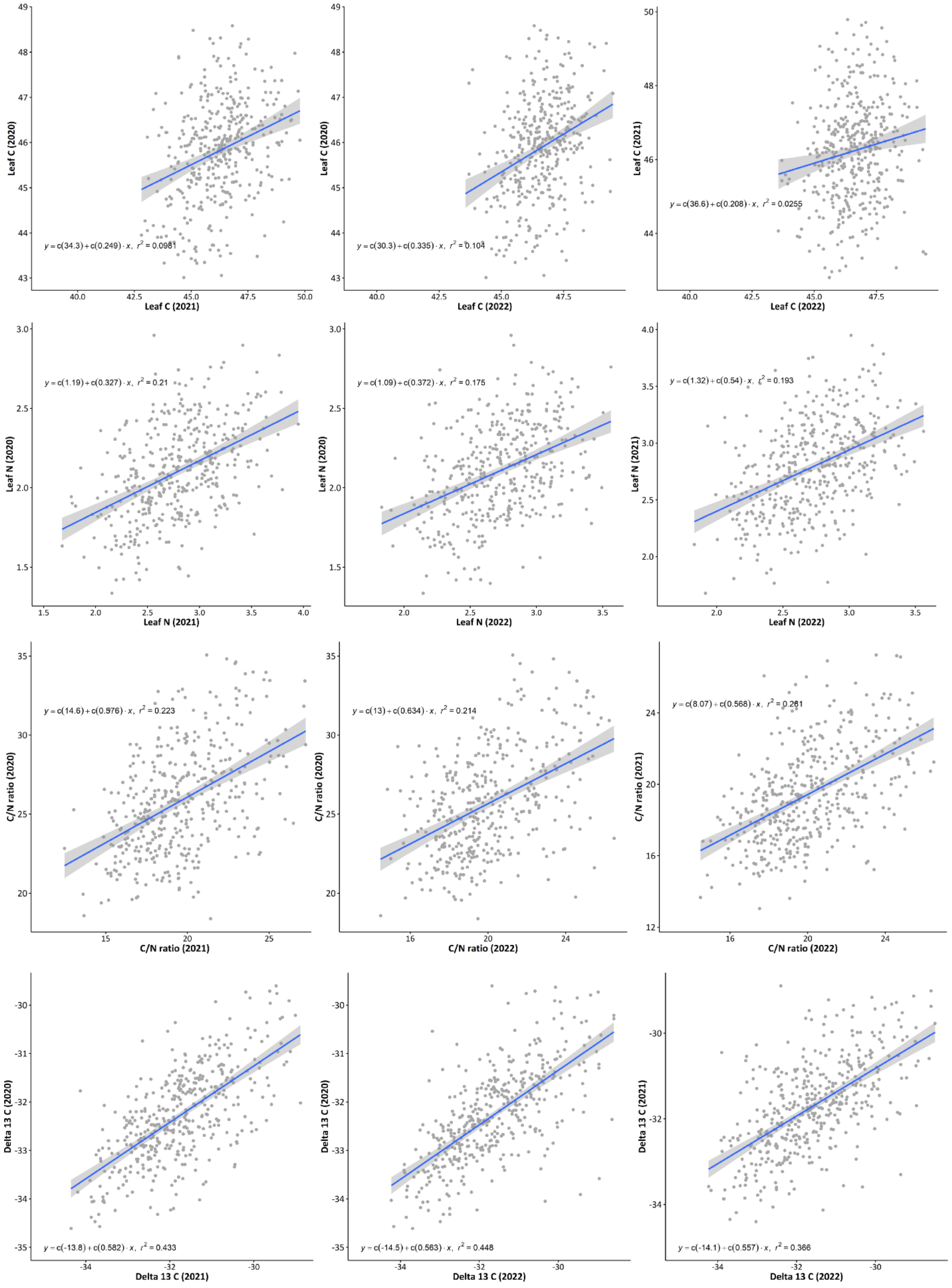

**S4:** Average monthly temperature in °C and monthly precipitation (mm) measured during the sampling season from April to September in 2020, 2021 and 2022 in Ruia (a), Lupului (b), Solomon (c), Tampa (d) and Lempes (e), which were used in the environmental principal component analysis (PCA).

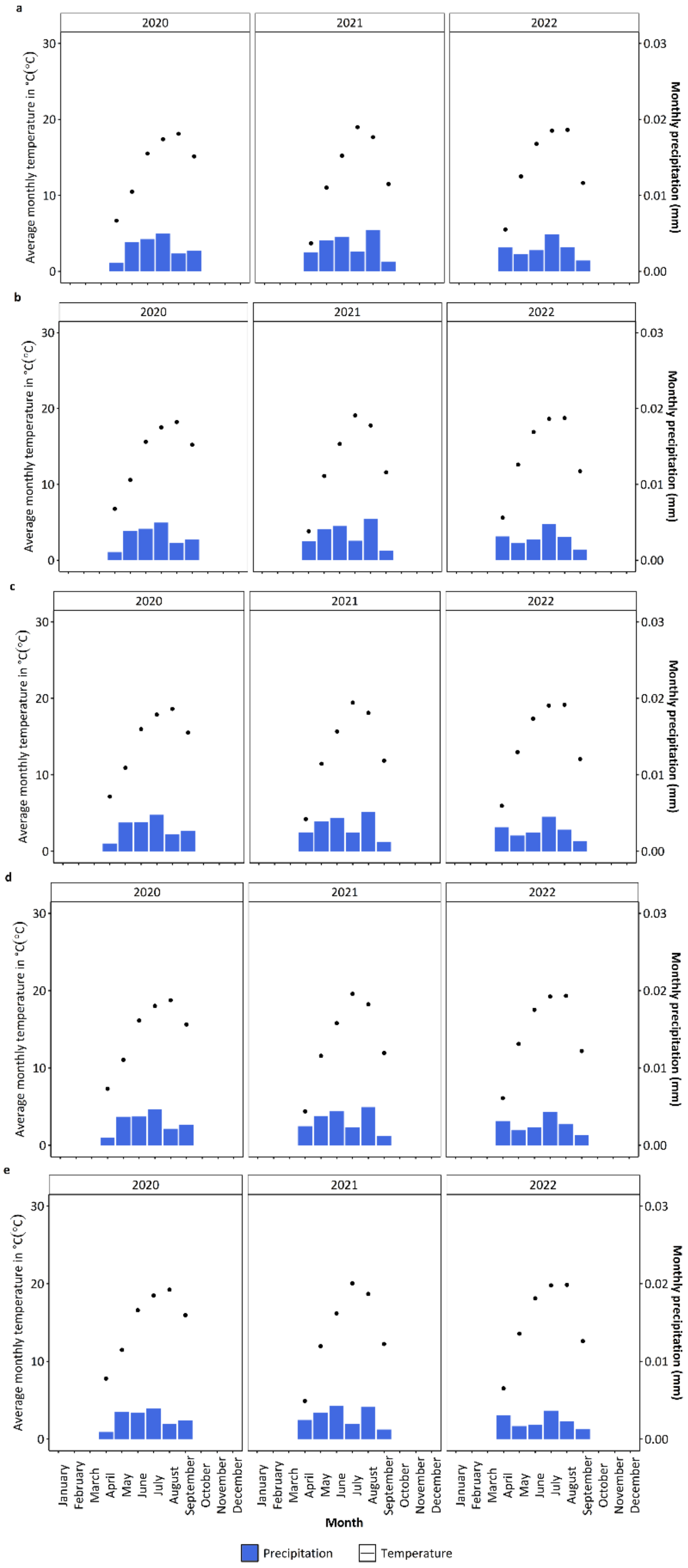

**S5:** Average monthly temperature in °C and monthly precipitation (mm) measured during the sampling season from April to September in 2020, 2021 and 2022 in Ruia, Lupului, Solomon, Tampa and Lempes, which were used in the environmental principal component analysis (PCA).

| Stand | Year | Stand | Precipitation in mm | Temperature in °C |
|---|---|---|---|---|
| April | 2020 | Ruia | 0.001097 | 6.68 |
| April | 2020 | Lupului | 0.001082 | 6.79 |
| April | 2020 | Solomon | 0.001027 | 7.16 |
| April | 2020 | Tampa | 0.001003 | 7.32 |
| April | 2020 | Lempes | 0.000903 | 7.79 |
| May | 2020 | Ruia | 0.003846 | 10.49 |
| May | 2020 | Lupului | 0.003833 | 10.58 |
| May | 2020 | Solomon | 0.003749 | 10.92 |
| May | 2020 | Tampa | 0.00369 | 11.06 |
| May | 2020 | Lempes | 0.003501 | 11.48 |
| June | 2020 | Ruia | 0.004246 | 15.52 |
| June | 2020 | Lupului | 0.004148 | 15.62 |
| June | 2020 | Solomon | 0.003832 | 15.98 |
| June | 2020 | Tampa | 0.00372 | 16.14 |
| June | 2020 | Lempes | 0.00336 | 16.60 |
| July | 2020 | Ruia | 0.004998 | 17.40 |
| July | 2020 | Lupului | 0.004955 | 17.51 |
| July | 2020 | Solomon | 0.004757 | 17.88 |
| July | 2020 | Tampa | 0.004615 | 18.03 |
| July | 2020 | Lempes | 0.003897 | 18.48 |
| August | 2020 | Ruia | 0.002331 | 18.11 |
| August | 2020 | Lupului | 0.002303 | 18.23 |
| August | 2020 | Solomon | 0.002197 | 18.62 |
| August | 2020 | Tampa | 0.002146 | 18.77 |
| August | 2020 | Lempes | 0.001981 | 19.23 |
| April | 2021 | Ruia | 0.002523 | 3.70 |
| April | 2021 | Lupului | 0.002513 | 3.82 |
| April | 2021 | Solomon | 0.002485 | 4.22 |
| April | 2021 | Tampa | 0.002478 | 4.39 |
| April | 2021 | Lempes | 0.002451 | 4.91 |
| May | 2021 | Ruia | 0.004061 | 11.02 |
| May | 2021 | Lupului | 0.004053 | 11.11 |
| May | 2021 | Solomon | 0.003926 | 11.44 |
| May | 2021 | Tampa | 0.003799 | 11.57 |
| May | 2021 | Lempes | 0.003399 | 11.95 |
| June | 2021 | Ruia | 0.00456 | 15.22 |
| June | 2021 | Lupului | 0.0045 | 15.33 |
| June | 2021 | Solomon | 0.004386 | 15.67 |
| June | 2021 | Tampa | 0.004396 | 15.79 |
| June | 2021 | Lempes | 0.004276 | 16.18 |
| July | 2021 | Ruia | 0.002571 | 18.99 |
| July | 2021 | Lupului | 0.002556 | 19.09 |
| July | 2021 | Solomon | 0.002439 | 19.44 |
| July | 2021 | Tampa | 0.002333 | 19.59 |
| July | 2021 | Lempes | 0.001943 | 20.05 |
| August | 2021 | Ruia | 0.005456 | 17.67 |
| August | 2021 | Lupului | 0.005416 | 17.77 |
| August | 2021 | Solomon | 0.005146 | 18.10 |
| August | 2021 | Tampa | 0.004929 | 18.24 |
| August | 2021 | Lempes | 0.004124 | 18.68 |
| April | 2022 | Ruia | 0.003118 | 5.52 |
| April | 2022 | Lupului | 0.003131 | 5.61 |
| April | 2022 | Solomon | 0.003158 | 5.95 |
| April | 2022 | Tampa | 0.003144 | 6.10 |
| April | 2022 | Lempes | 0.003019 | 6.53 |
| May | 2022 | Ruia | 0.002278 | 12.50 |

| | | | | |
|---|---|---|---|---|
| May | 2022 | Lupului | 0.002242 | 12.61 |
| May | 2022 | Solomon | 0.002058 | 12.98 |
| May | 2022 | Tampa | 0.001948 | 13.12 |
| May | 2022 | Lempes | 0.001672 | 13.57 |
| June | 2022 | Ruia | 0.002778 | 16.78 |
| June | 2022 | Lupului | 0.002709 | 16.91 |
| June | 2022 | Solomon | 0.002456 | 17.34 |
| June | 2022 | Tampa | 0.002332 | 17.53 |
| June | 2022 | Lempes | 0.00181 | 18.12 |
| July | 2022 | Ruia | 0.004829 | 18.52 |
| July | 2022 | Lupului | 0.004786 | 18.64 |
| July | 2022 | Solomon | 0.004518 | 19.06 |
| July | 2022 | Tampa | 0.004322 | 19.24 |
| July | 2022 | Lempes | 0.003657 | 19.78 |
| August | 2022 | Ruia | 0.003133 | 18.63 |
| August | 2022 | Lupului | 0.003066 | 18.76 |
| August | 2022 | Solomon | 0.002836 | 19.16 |
| August | 2022 | Tampa | 0.002729 | 19.33 |
| August | 2022 | Lempes | 0.002308 | 19.85 |

**S6:** Results from the principal coordinate analysis (PCoA) on precipitation and temperature, of the sampling years 2020 (**a**) and the eigenvalues of PCs (**b**), 2021 (**c** and **d**) and 2022 (**e** and **f**). Environmental PC 1, which explained ~100% was used in the genome-wide association study (GWAS) as covariate.

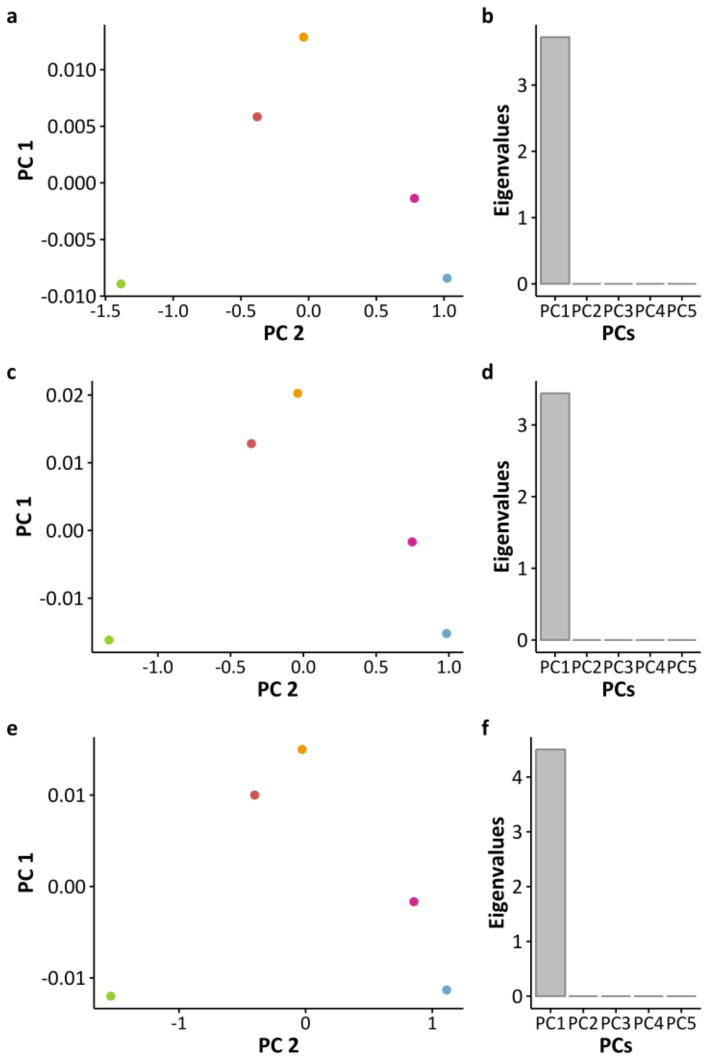

**S7:** QQ-Plots of all phenotypic variables. The QQ plots are following this order: Stomatal density (**a**), $\delta^{13}C$ measured in 2020 (**b**), $\delta^{13}C$ measured in 2021 (**c**), $\delta^{13}C$ measured in 2022 (d), leaf nitrogen measured in 2020 (**e**), leaf nitrogen measured in 2021 (**f**), leaf nitrogen measured in 2022 (**g**), C/N ratio in 2020 (**h**), C/N ratio in 2021 (**i**), C/N ratio in 2022 (**j**).

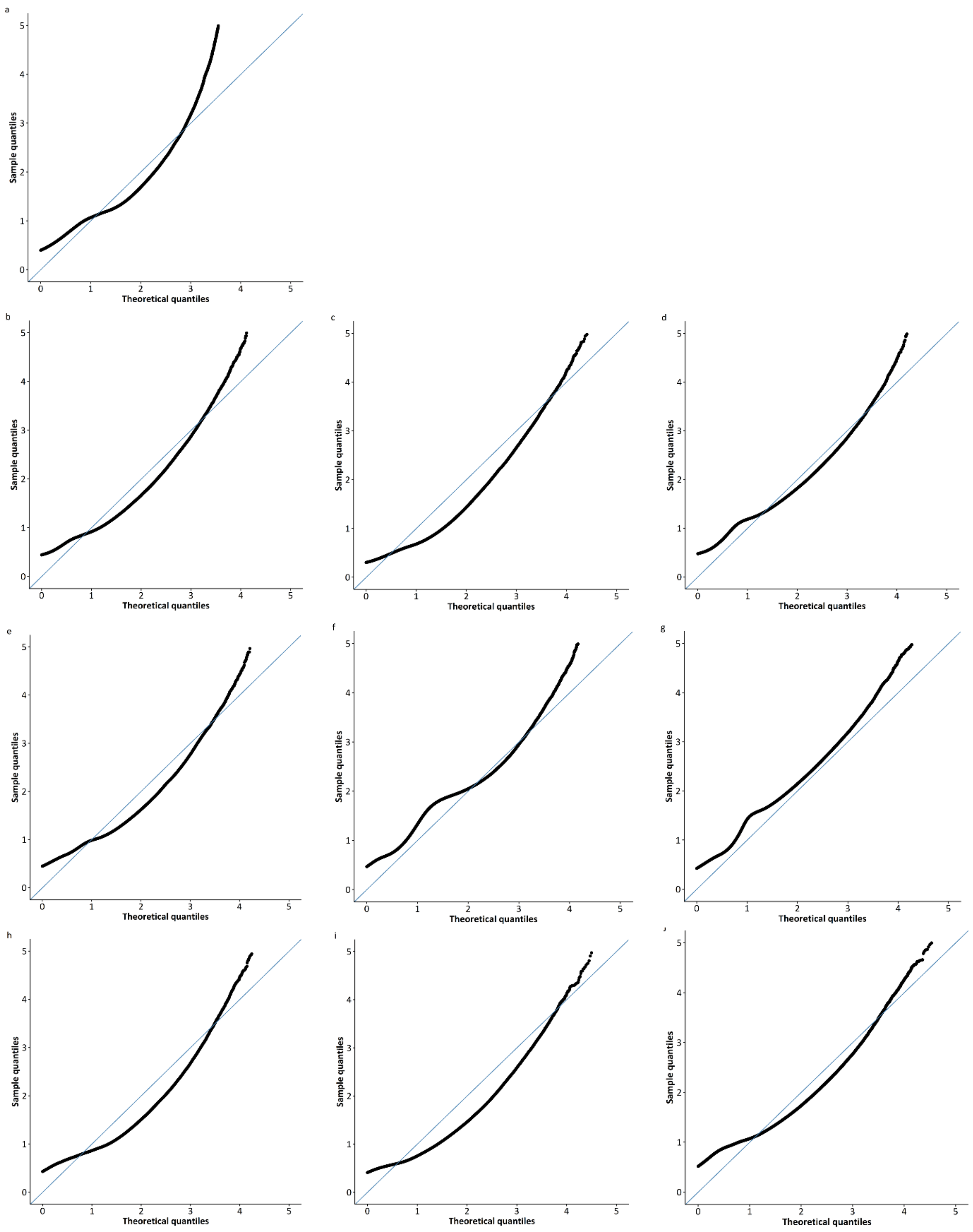


**S8:** List of remaining candidate SNPs associated with stomatal density (SD) and $\delta^{13}C$ measured in 2020 with premutation derived significance levels of p ≤0.000001 and their annotated gene variants, the underlying genes with their locations, functions and predicted impact (LOW or MODIFIER (not predicted)) based on SnpEff (Cingolani et al. 2012).

| Trait | Chr | Position of SNP | Distance to gene variant in kb | Gene variant | Gene |
|---|---|---|---|---|---|
| SD | 4 | 40808183 | NA | NA | NA |
| | | 40808384 | NA | NA | NA |
| | | 40816400 | NA | NA | NA |
| | | 40818194 | NA | NA | NA |
| | | 40854731 | NA | NA | NA |
| | 5 | 40871447 | NA | NA | NA |
| | 6 | 15825445 | NA | NA | NA |
| | | 15825457 | NA | NA | NA |
| | | 15825492 | NA | NA | 15825492 |
| | | 15825793 | NA | NA | 15825793 |
| | | 15825811 | NA | NA | 15825811 |
| | 8 | 7588499 | 2.522 | Bhaga_8.g971 | NA |
| | | | 2.505 | Bhaga_8.g971 | NA |
| | | | 2.206 | Bhaga_8.g970 | NA |
| | 9 | 14893568 | ~-6.168 to -6.31 | Bhaga_9.g1841 | TYK12172.1 |
| | | | ~6.161 | Bhaga_9.g1841 | TYK12172.1 |
| | | | ~6.207 | Bhaga_9.g1841 | TYK12172.1 |
| | | 5470445 | NA | NA | NA |
| | | 6025501 | NA | NA | NA |
| | | 6851324 | NA | NA | NA |
| | | 6858935 | NA | NA | NA |
| | | 6876799 | NA | NA | NA |
| | | 6970293 | NA | NA | NA |
| | | 7116359 | ~-6.0 | Bhaga_10.g853 | KAB1223135.1 |
| | | 7116359 | ~-7.8 | Bhaga_10.g853 | KAB1223135.1 |
| | | 7116359 | ~-14.2 | Bhaga_10.g857 | XP_018814658.1 |
| | | 7211779 | ~-9.1 to 14 | Bhaga_10.g871 | XP_023888712.1 |
| | | 7513203 | NA | NA | NA |
| | | 7776590 | NA | NA | NA |
| | | 7787247 | NA | NA | NA |
| | | 7825443 | NA | NA | NA |
| | | 8260485 | NA | NA | NA |
| | | 8260499 | NA | NA | NA |
| | | 8323901 | NA | NA | NA |
| | | 8399501 | NA | NA | NA |
| | | 8586058 | NA | NA | NA |
| | | 9215197 | NA | NA | NA |
| | | 9520963 | NA | NA | NA |
| | | 8586058 | NA | NA | NA |
| | | 9215197 | NA | NA | NA |
| | | 9520963 | NA | NA | NA |
| | | 10058669 | NA | NA | NA |
| | | 10159262 | NA | NA | NA |
| | | 10178899 | NA | NA | NA |
| | | 10527710 | NA | NA | NA |
| | | 10540324 | NA | NA | NA |
| | | 10644942 | NA | NA | NA |
| | | 10644983 | NA | NA | NA |
| | | 10653218 | NA | NA | NA |
| | | 10688216 | NA | NA | NA |
| | | 10806739 | NA | NA | NA |

| | | | | |
|---|---|---|---|---|
| | 10822774 | NA | NA | NA |
| $\delta^{13}C$ (2020) | 10907107 | -3.796 | Bhaga_10.g1348 | XP_030940353.1 |
| | 10907267 | -3.643 | Bhaga_10.g1348 | |
| SD | 10909086 | -1.853 | Bhaga_10.g1353 | |
| | 10909126 | ~-18.6 to -18.4 | Bhaga_10.g1353 | PQQ02746.1 |
| | 10909168 | | | |
| | 10909306 | | | |
| | 10909343 | | | |
| | 10915528 | 2.844 | Bhaga_10.g1350 | XP_030941044.1 |
| | 11233527 | ~-9.6 to -11.4 | Bhaga_10.g1390 | XP_023922841.1 |
| | 11233527 | | | |
| | 11233581 | | | |
| | 11401610 | NA | NA | NA |
| | 11467008 | NA | NA | NA |
| | 11467225 | NA | NA | NA |
| | 11511420 | ~6.9 to 10.4 | Bhaga_10.g1422 | XP_023884902.1 |
| | | ~-11.4 | Bhaga_10.g1424 | KAB1212694.1 |
| | 11599598 | NA | NA | NA |
| | 11634927 | NA | NA | NA |
| | 11988800 | NA | NA | NA |
| | 12053423 | NA | NA | NA |
| | 12684288 | NA | NA | NA |

**S9:** Minor allele frequency (MAF) observed in the different stands at the significant markers (p ≤0.000001) on chromosome 10 associated with stomatal density and the number of heterozygous individuals (Nr of Het), number of homozygous individuals (Nr of Hom) and the total number of genotyped individuals (Nr of total ind.).

| Marker | Stand | MAF | Nr of Het. | Nr of Hom. | Nr of total ind. | Marker | MAF | Nr of Het | Nr of Hom | Nr of total ind. |
|---|---|---|---|---|---|---|---|---|---|---|
| chr10_1090912 | Lempes | 0 | 0 | 0 | 95 | chr10_1367781 | 0 | 0 | 0 | 67 |
| chr10_1090912 | Tampa | 0 | 0 | 0 | 85 | chr10_1367781 | 0 | 0 | 0 | 56 |
| chr10_1090912 | Solomon | 0.0217 | 4 | 0 | 92 | chr10_1367781 | 0.0085 | 1 | 0 | 59 |
| chr10_1090912 | Lupului | 0.1312 | 19 | 1 | 80 | chr10_1367781 | 0.0625 | 6 | 0 | 48 |
| chr10_1090912 | Ruia | 0.4899 | 65 | 16 | 99 | chr10_1367781 | 0.3661 | 13 | 14 | 56 |
| chr10_1090916 | Lempes | 0 | 0 | 0 | 95 | chr10_489837 | 0.02 | 4 | 0 | 100 |
| chr10_1090916 | Tampa | 0 | 0 | 0 | 91 | chr10_489837 | 0.0051 | 1 | 0 | 99 |
| chr10_1090916 | Solomon | 0.0204 | 4 | 0 | 98 | chr10_489837 | 0.0102 | 2 | 0 | 98 |
| chr10_1090916 | Lupului | 0.1359 | 21 | 2 | 92 | chr10_489837 | 0.1061 | 19 | 1 | 99 |
| chr10_1090916 | Ruia | 0.4848 | 64 | 16 | 99 | chr10_489837 | 0.2879 | 45 | 6 | 99 |
| chr10_1090930 | Lempes | 0 | 0 | 0 | 100 | chr10_631415 | 0 | 0 | 0 | 100 |
| chr10_1090930 | Tampa | 0 | 0 | 0 | 99 | chr10_631415 | 0 | 0 | 0 | 99 |
| chr10_1090930 | Solomon | 0.0204 | 4 | 0 | 98 | chr10_631415 | 0.0102 | 2 | 0 | 98 |
| chr10_1090930 | Lupului | 0.1224 | 22 | 1 | 98 | chr10_631415 | 0.096 | 19 | 0 | 99 |
| chr10_1090930 | Ruia | 0.4798 | 65 | 15 | 99 | chr10_631415 | 0.3485 | 53 | 8 | 99 |
| chr10_1090934 | Lempes | 0 | 0 | 0 | 100 | chr10_711635 | 0.005 | 1 | 0 | 100 |
| chr10_1090934 | Tampa | 0 | 0 | 0 | 99 | chr10_711635 | 0.0051 | 1 | 0 | 99 |
| chr10_1090934 | Solomon | 0.0204 | 4 | 0 | 98 | chr10_711635 | 0.0306 | 4 | 1 | 98 |
| chr10_1090934 | Lupului | 0.1224 | 22 | 1 | 98 | chr10_711635 | 0.1061 | 17 | 2 | 99 |
| chr10_1090934 | Ruia | 0.4798 | 65 | 15 | 99 | chr10_711635 | 0.3838 | 50 | 13 | 99 |
| chr10_1123352 | Lempes | 0.0054 | 1 | 0 | 93 | chr10_719880 | 0.015 | 3 | 0 | 100 |
| chr10_1123352 | Tampa | 0 | 0 | 0 | 97 | chr10_719880 | 0.0354 | 7 | 0 | 99 |
| chr10_1123352 | Solomon | 0.0215 | 2 | 1 | 93 | chr10_719880 | 0.0357 | 7 | 0 | 98 |
| chr10_1123352 | Lupului | 0.1276 | 21 | 2 | 98 | chr10_719880 | 0.1364 | 27 | 0 | 99 |
| chr10_1123352 | Ruia | 0.5808 | 45 | 35 | 99 | chr10_719880 | 0.3838 | 52 | 12 | 99 |
| chr10_1123358 | Lempes | 0.0053 | 1 | 0 | 94 | chr10_721177 | 0.0052 | 1 | 0 | 96 |
| chr10_1123358 | Tampa | 0 | 0 | 0 | 97 | chr10_721177 | 0.0152 | 3 | 0 | 99 |
| chr10_1123358 | Solomon | 0.0215 | 2 | 1 | 93 | chr10_721177 | 0.0204 | 4 | 0 | 98 |
| chr10_1123358 | Lupului | 0.1276 | 21 | 2 | 98 | chr10_721177 | 0.1042 | 20 | 0 | 96 |

| | | | | | | | | | |
|---|---|---|---|---|---|---|---|---|---|
| chr10_1123358 | Ruia | 0.5808 | 45 | 35 | 99 | chr10_721177 | 0.3687 | 51 | 11 | 99 |
| chr10_1151142 | Lempes | 0.0643 | 1 | 4 | 70 | chr10_755543 | 0 | 0 | 0 | 99 |
| chr10_1151142 | Tampa | 0.037 | 0 | 2 | 54 | chr10_755543 | 0.0253 | 5 | 0 | 99 |
| chr10_1151142 | Solomon | 0.058 | 2 | 3 | 69 | chr10_755543 | 0.0255 | 5 | 0 | 98 |
| chr10_1151142 | Lupului | 0.2041 | 4 | 8 | 49 | chr10_755543 | 0.1031 | 20 | 0 | 97 |
| chr10_1151142 | Ruia | 0.7011 | 12 | 55 | 87 | chr10_755543 | 0.4343 | 54 | 16 | 99 |
| chr10_1190395 | Lempes | 0.0165 | 3 | 0 | 91 | chr10_778592 | 0 | 0 | 0 | 100 |
| chr10_1190395 | Tampa | 0.0054 | 1 | 0 | 93 | chr10_778592 | 0.0101 | 2 | 0 | 99 |
| chr10_1190395 | Solomon | 0.0054 | 1 | 0 | 92 | chr10_778592 | 0.0102 | 2 | 0 | 98 |
| chr10_1190395 | Lupului | 0.1292 | 21 | 1 | 89 | chr10_778592 | 0.102 | 20 | 0 | 98 |
| chr10_1190395 | Ruia | 0.5312 | 56 | 23 | 96 | chr10_778592 | 0.4242 | 56 | 14 | 99 |
| chr10_1254710 | Lempes | 0.0104 | 2 | 0 | 96 | chr10_778597 | 0 | 0 | 0 | 100 |
| chr10_1254710 | Tampa | 0.0206 | 4 | 0 | 97 | chr10_778597 | 0.0102 | 2 | 0 | 98 |
| chr10_1254710 | Solomon | 0.0206 | 4 | 0 | 97 | chr10_778597 | 0.0102 | 2 | 0 | 98 |
| chr10_1254710 | Lupului | 0.1141 | 17 | 2 | 92 | chr10_778597 | 0.1031 | 20 | 0 | 97 |
| chr10_1254710 | Ruia | 0.4394 | 57 | 15 | 99 | chr10_778597 | 0.4242 | 56 | 14 | 99 |
| chr10_1266723 | Lempes | 0.064 | 9 | 1 | 86 | chr10_778746 | 0 | 0 | 0 | 96 |
| chr10_1266723 | Tampa | 0.0959 | 10 | 2 | 73 | chr10_778746 | 0.0102 | 2 | 0 | 98 |
| chr10_1266723 | Solomon | 0.0769 | 12 | 0 | 78 | chr10_778746 | 0.0103 | 2 | 0 | 97 |
| chr10_1266723 | Lupului | 0.189 | 27 | 2 | 82 | chr10_778746 | 0.1064 | 20 | 0 | 94 |
| chr10_1266723 | Ruia | 0.3646 | 66 | 2 | 96 | chr10_778746 | 0.4242 | 56 | 14 | 99 |
| chr10_1343528 | Lempes | 0 | 0 | 0 | 72 | chr10_782027 | 0 | 0 | 0 | 84 |
| chr10_1343528 | Tampa | 0.0204 | 2 | 0 | 49 | chr10_782027 | 0.0128 | 1 | 0 | 39 |
| chr10_1343528 | Solomon | 0.0211 | 3 | 0 | 71 | chr10_782027 | 0.0068 | 1 | 0 | 74 |
| chr10_1343528 | Lupului | 0.1071 | 7 | 1 | 42 | chr10_782027 | 0.0714 | 6 | 0 | 42 |
| chr10_1343528 | Ruia | 0.2722 | 31 | 6 | 79 | chr10_782027 | 0.4026 | 42 | 10 | 77 |
| chr10_1367775 | Lempes | 0 | 0 | 0 | 64 | chr10_782038 | 0.0101 | 2 | 0 | 99 |
| chr10_1367775 | Tampa | 0 | 0 | 0 | 55 | chr10_782038 | 0.0532 | 10 | 0 | 94 |
| chr10_1367775 | Solomon | 0.0086 | 1 | 0 | 58 | chr10_782038 | 0.0412 | 8 | 0 | 97 |
| chr10_1367775 | Lupului | 0.0581 | 5 | 0 | 43 | chr10_782038 | 0.1474 | 26 | 1 | 95 |
| chr10_1367775 | Ruia | 0.3725 | 8 | 15 | 51 | chr10_782038 | 0.4286 | 56 | 14 | 98 |

**S10:** Correlation between stomatal density and the minor allele frequency (MAF) of the significant markers (p ≤0.000001) on chromosome 10 at ~ 6.314 at 0.92 (p-value = 0.028) (a), ~ 7.116 at 0.9 (p-value = 0.037) (b), ~ 7.555 at 0.89 (p-value = 0.041) (c), ~ 7.7859 at 0.91 (p-value = 0.033) (d), ~ 7.82 at 0.89 (p-value = 0.045) (e), ~ 10.90916 at 0.92 (p-value = 0.029) (f), ~ 11.2335 at 0.9 (p-value = 0.039) (g), ~ 12.5471 at 0.91 (p-value = 0.033) (h), ~ 13.67776 at 0.88 (p-value = 0.047) (i).

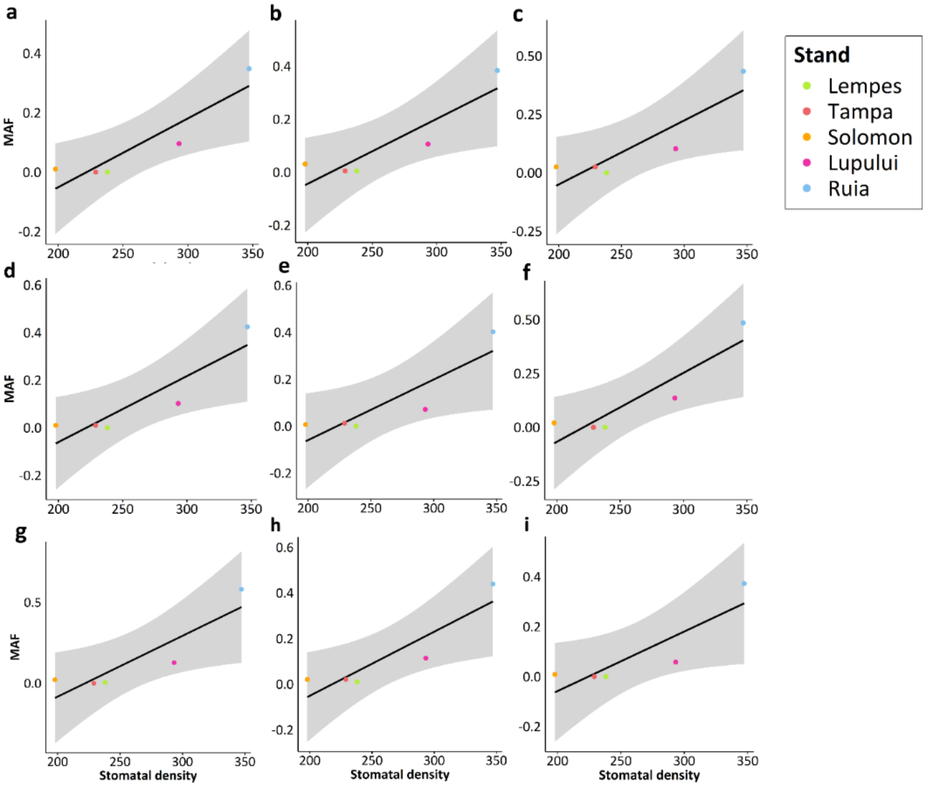


**S11:** Minor allele frequency (MAF) observed in the different stands at the significant markers (p ≤0.000001) associated with leaf nitrogen content and C/N ratio, the stand mean, the number of heterozygous individuals (Nr of Het), number of homozygous individuals (Nr of Hom) and the total number of genotyped individuals (Nr of total ind.).

| **Marker** | **Population** | **MAF** | **Nr of Het.** | **Nr of Hom.** | **Nr of total** | **Trait** | **Trait mean** |
|---|---|---|---|---|---|---|---|
| chr2_31803661 | Lempes | 0.5 | 41 | 26 | 93 | NC_2021 | 18.39 |
| chr2_31803661 | Tampa | 0.401042 | 45 | 16 | 96 | NC_2021 | 19.06 |
| chr2_31803661 | Solomon | 0.380208 | 39 | 17 | 96 | NC_2021 | 21.45 |
| chr2_31803661 | Lupului | 0.406593 | 54 | 10 | 91 | NC_2021 | 20.53 |
| chr2_31803661 | Ruia | 0.405263 | 51 | 13 | 95 | NC_2021 | 18.92 |
| chr10_13027242 | Lempes | 0.06707 | 7 | 2 | 82 | CNR 2020 | 2.09 |
| chr10_13027242 | Tampa | 0.05172 | 7 | 1 | 87 | CNR 2020 | 2.08 |
| chr10_13027242 | Solomon | 0.10440 | 11 | 4 | 91 | CNR 2020 | 2.01 |
| chr10_13027242 | Lupului | 0.11702 | 14 | 4 | 94 | CNR 2020 | 2.21 |
| chr10_13027242 | Ruia | 0.02604 | 3 | 1 | 96 | CNR 2020 | 2.20 |
| chr11_26539169 | Lempes | 0.13 | 22 | 2 | 100 | NC_2020 | 25.70 |
| chr11_26539169 | Tampa | 0.131313 | 26 | 0 | 99 | NC_2020 | 25.59 |
| chr11_26539169 | Solomon | 0.086735 | 17 | 0 | 98 | NC_2020 | 27.36 |
| chr11_26539169 | Lupului | 0.163265 | 24 | 4 | 98 | NC_2020 | 25.49 |
| chr11_26539169 | Ruia | 0.176768 | 23 | 6 | 99 | NC_2020 | 25.26 |

**S12:** Correlation between leaf nitrogen content measured in 2021 and the minor allele frequency (MAF) of the significant marker (p ≤0.000001) on chromosome 2 at ~31.804 Mb. This marker was annotated with the gene variant *Bhaga_2.g3457* with the underlying gene *XP_030960280.1*, also known as *abscisic-aldehyde oxidase-like*.

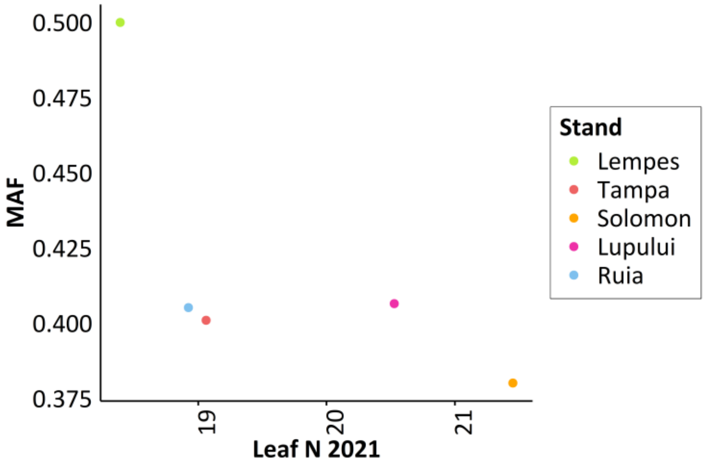


**S13:** Correlation between C/N ratio measured in 2020 and the minor allele frequency (MAF) of the significant marker (p ≤0.000001) on chromosome 10 at ~13.03 Mb. No annotations were found for this marker.

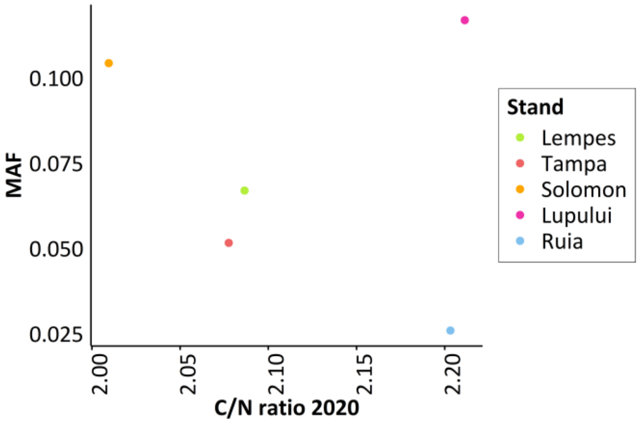


**S14:** Rate of false positive observations observed in Ghat test calculated based on the 100 randomly generated traits with 10 replications (10 replications x 100 random traits).

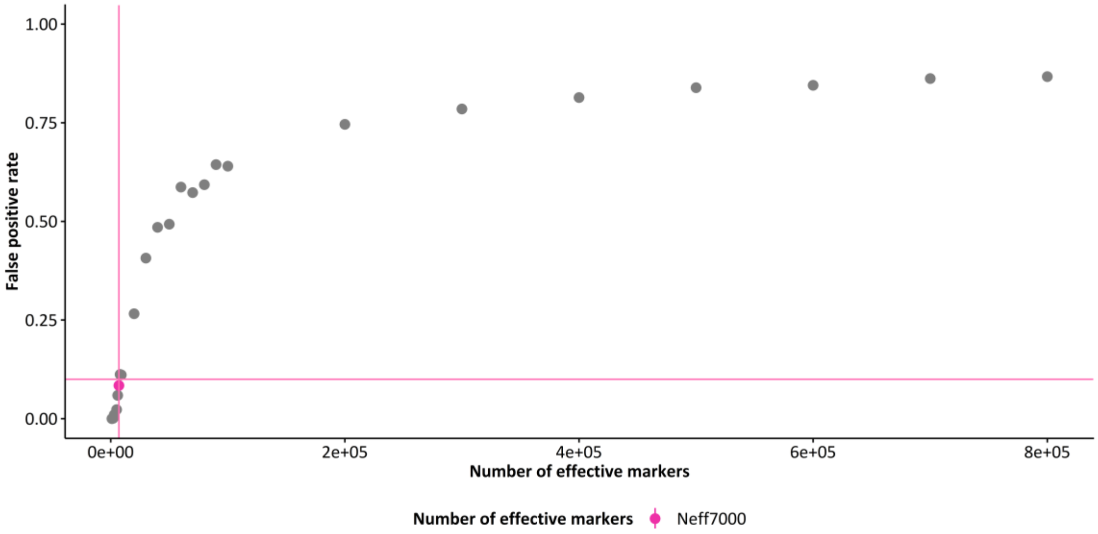

**S15:** Results from five-fold cross-validation with the mean predictive ability as $r_{y\hat{g}}$ (correlation of predicted trait estimates and observed trait measurements) with random partition of stands in test and training set (**a**) and five-fold cross-validation in which one stand was left out and the remaining stands were used for prediction (**b**) with 100 replications based on RR-BLUP for the real trait measurements.

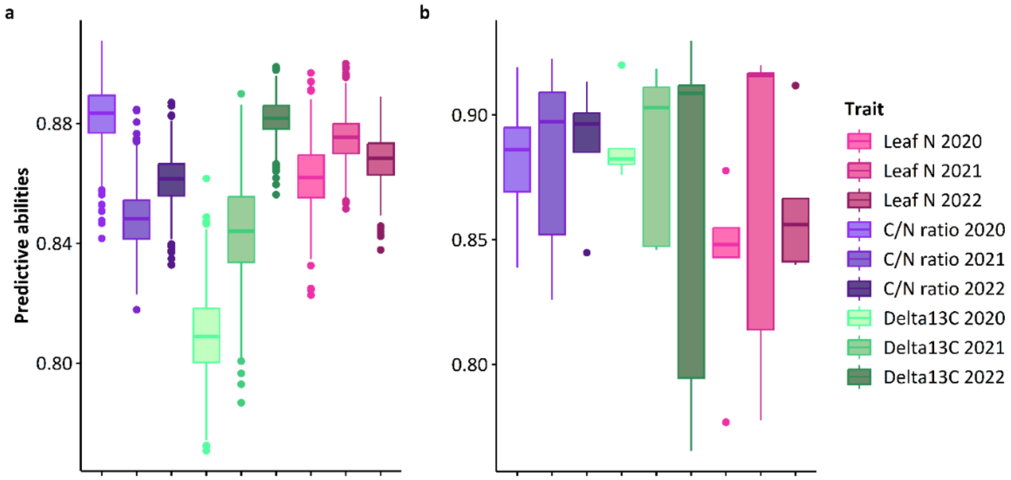


**S16:** Results from $\hat{G}$ test with change calculated between groups (Ruia, Solomon vs Tampa, Lempes) calculated between groups with 100000 permutations and 7000 effective markers calculated based on the previous test runs for carbon isotope composition $\delta^{13}C$.

| Trait | $\hat{G}$ correlation | p-value |
|---|---|---|
| $\delta^{13}C$ (2020) | 0.1017 | 0.0000 |
| $\delta^{13}C$ (2021) | 0.0962 | 0.0000 |
| $\delta^{13}C$ (2022) | -0.017 | 0.1527 |

**S17:** Coding region of the gene variant *Bhaga_8.g1863* (*ABC transporter I family member 10*) and the significant marker on chromosome 8 at ~ 15.485 Mb in blue and pairwise LD heat map from 15470128 to 15500213 Mb.

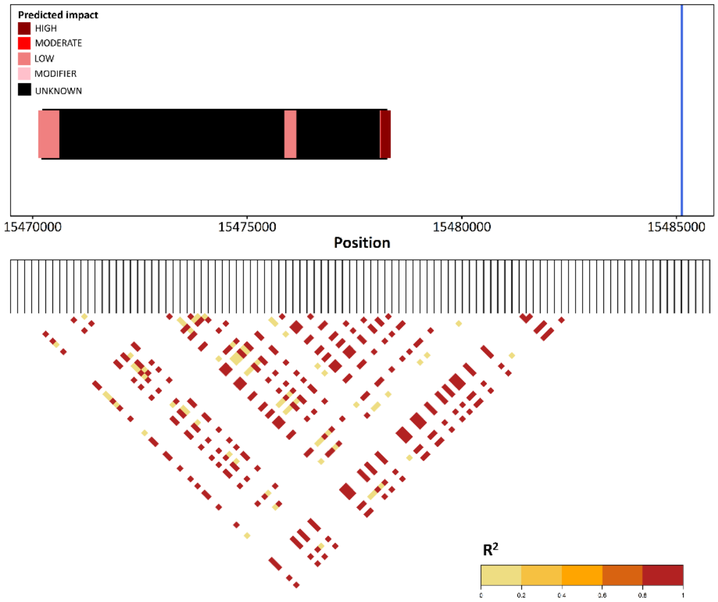

**S18:** Coding region of the gene variant *Bhaga_10.g917* (*ALMT2*) and the significant marker on chromosome 10 at ~7.56 Mb in blue and pairwise LD heat map from 7.550433 to 7560433 Mb.

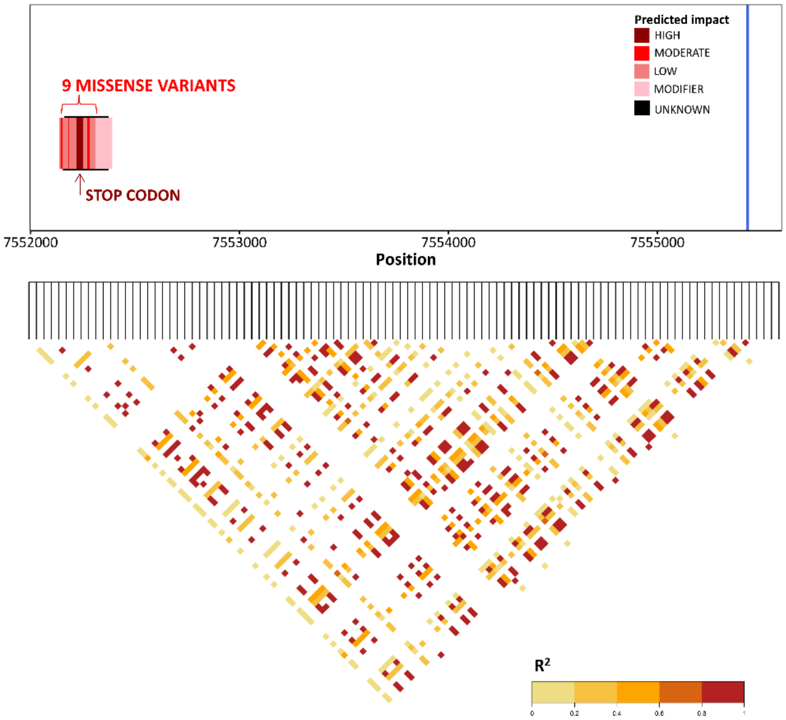


**S19:** Observed codon changes which lead to an amino acid (AA) replacement and a resulting change in polar bonds (hydrophobic or hydrophilic) at the underlying gene variants of markers associated with stomatal density

| Gene variant | Variant version | Change in polar bonds | Original | Replaced by |
|---|---|---|---|---|
| Bhaga_8.g506 | 5 missense variants | Hydrophobic AA was replaced by hydrophilic AA | Pro | Ser |
| | | Hydrophilic AA was replaced by hydrophobic AA | Glu | Asp |
| | | Hydrophilic AA was replaced by hydrophobic AA | Arg | Lys |
| | | Hydrophilic AA was replaced by hydrophobic AA | Val | Ala |
| | | Hydrophobic AA was replaced by hydrophilic AA | Asn | Lys |
| | | Hydrophilic AA was replaced by hydrophobic AA | Glu | Asp |
| | 3 missense variants | no change in polar bonds | | |
| Bhaga_8.g1863 | 1 missense variant | Hydrophilic AA was replaced by hydrophobic AA | Lys | Asn |
| Bhaga_9.g1680 | 1 missense variant | Hydrophilic AA was replaced by hydrophobic AA | Asp | His |
| Bhaga_9.g1682 | 13 missense variants | Hydrophobic AA was replaced by hydrophilic AA | Asn | Lys |
| | | Hydrophilic AA was replaced by hydrophobic AA | Arg | Tyr |
| | | Hydrophilic AA was replaced by hydrophobic AA | Asn | Ile |
| | | Hydrophilic AA was replaced by hydrophobic AA | Lys | Glu |
| | | Hydrophilic AA was replaced by hydrophobic AA | Glu | Asp |
| | | Hydrophilic AA was replaced by hydrophobic AA | Thr | Ser |
| | | Hydrophobic AA was replaced by hydrophilic AA | Tyr | Asp |
| | | Hydrophilic AA was replaced by hydrophobic AA | Lys | Glu |
| | | Hydrophilic AA was replaced by hydrophobic AA | Arg | Gly |
| | | Hydrophobic AA was replaced by hydrophilic AA | Gly | Asp |
| | | Hydrophobic AA was replaced by hydrophilic AA | Asn | Ser |
| | | Hydrophilic AA was replaced by hydrophobic AA | His | Leu |
| | | Hydrophilic AA was replaced by hydrophobic AA | Gly | Val |
| Bhaga_10.g591 | 3 missense variants | Hydrophobic AA was replaced by hydrophilic AA | Leu | His |
| | | Hydrophobic AA was replaced by hydrophilic AA | Asn | His |
| | | Hydrophilic AA was replaced by hydrophobic AA | Leu | Val |
| Bhaga_10.g752 | 1 missense variant | Hydrophilic AA was replaced by hydrophobic AA | Glu | Lys |
| Bhaga_10.g753 | 3 missense variants | Hydrophilic AA was replaced by hydrophobic AA | Val | Ile |

| | | | | |
|---|---|---|---|---|
| | | no change in polar bonds | Gln | Lys |
| | | Hydrophobic AA was replaced by hydrophilic AA | Met | Lys |
| Bhaga_10.g852 | 1 missense variant | Hydrophilic AA was replaced by hydrophobic AA | Val | Ile |
| Bhaga_10.g854 | 7 missense variants | Hydrophilic AA was replaced by hydrophobic AA | Phe | Tyr |
| | | Hydrophilic AA was replaced by hydrophobic AA | Leu | Phe |
| | | Hydrophilic AA was replaced by hydrophobic AA | Ala | Val |
| | | Hydrophobic AA was replaced by hydrophilic AA | Asn | Lys |
| | | Hydrophilic AA was replaced by hydrophobic AA | Val | Ile |
| | | Hydrophilic AA was replaced by hydrophobic AA | Glu | Gly |
| | | Hydrophilic AA was replaced by hydrophobic AA | Phe | Val |
| Bhaga_10.g867 | 7 missense variants | Hydrophilic AA was replaced by hydrophobic AA | Thr | Asn |
| | | Hydrophilic AA was replaced by hydrophobic AA | Leu | Phe |
| | | Hydrophilic AA was replaced by hydrophobic AA | Gly | Ala |
| | | Hydrophilic AA was replaced by hydrophobic AA | His | Leu |
| | | Hydrophilic AA was replaced by hydrophobic AA | Ser | Trp |
| | | Hydrophobic AA was replaced by hydrophilic AA | Asn | His |
| | | Hydrophobic AA was replaced by hydrophilic AA | Asn | Lys |
| Bhaga_10.g917 | 9 missense variants | Hydrophilic AA was replaced by hydrophobic AA | Phe | Leu |
| | | Hydrophilic AA was replaced by hydrophobic AA | Asp | Ala |
| | | Hydrophilic AA was replaced by hydrophobic AA | Ser | Cys |
| | | Hydrophilic AA was replaced by hydrophobic AA | Leu | Phe |
| | | Hydrophobic AA was replaced by hydrophilic AA | Ile | Ser |
| | | Hydrophobic AA was replaced by hydrophilic AA | Gly | Glu |
| | | Hydrophobic AA was replaced by hydrophilic AA | Gly | Arg |
| | | Hydrophilic AA was replaced by hydrophobic AA | Ser | Leu |
| | | Hydrophilic AA was replaced by hydrophobic AA | Ser | Asn |
| Bhaga_10.g957 | 5 missense variants | Hydrophilic AA was replaced by hydrophobic AA | Glu | Lys |
| | | Hydrophobic AA was replaced by hydrophilic AA | Asn | Lys |
| | | Hydrophobic AA was replaced by hydrophilic AA | Pro | Thr |
| | | Hydrophilic AA was replaced by hydrophobic AA | Ala | Val |
| | | Hydrophilic AA was replaced by hydrophobic AA | Arg | Gln |
| | 25 missense variants | no change in polar bonds | | |
| Bhaga_10.g1460 | 2 missense variants | Hydrophilic AA was replaced by hydrophobic AA | Gly | Ala |
| | | Hydrophilic AA was replaced by hydrophobic AA | Ile | Val |
| | 1 missense variant | no change in polar bonds | | |
| Bhaga_10.g1462 | 5 missense variants | Hydrophobic AA was replaced by hydrophilic AA | Ala | Ser |
| | | Hydrophilic AA was replaced by hydrophobic AA | Thr | Lys |
| | | Hydrophobic AA was replaced by hydrophilic AA | Gly | Arg |
| | | Hydrophilic AA was replaced by hydrophobic AA | Asp | Asn |
| | | Hydrophilic AA was replaced by hydrophobic AA | Asp | Glu |
| Bhaga_10.g1543 | 3 missense variants | Hydrophobic AA was replaced by hydrophilic AA | Tyr | His |
| | | Hydrophobic AA was replaced by hydrophilic AA | Phe | Cys |
| | | Hydrophilic AA was replaced by hydrophobic AA | Ser | Thr |
| Bhaga_10.g1652 | 1 missense variant | Hydrophilic AA was replaced by hydrophobic AA | Glu | Lys |
| Bhaga_10.g1674 | 5 missense variants | Hydrophilic AA was replaced by hydrophobic AA | Ile | Leu |
| | | Hydrophilic AA was replaced by hydrophobic AA | Val | Ala |
| | | Hydrophilic AA was replaced by hydrophobic AA | Arg | Pro |
| | | Hydrophilic AA was replaced by hydrophobic AA | Arg | Gly |
| | | Hydrophobic AA was replaced by hydrophilic AA | Leu | His |

**S20:** Coding region of the gene variant *Bhaga_10.g957* (*MYB35-like*) and the five significant markers on chromosome 10 at ~7.78 to 7.82 Mb ( 7785927; 7785978; 7787468; 7820271; 7820389) Mb in blue and pairwise LD heat map from 7.765927 to 7.840389 Mb.

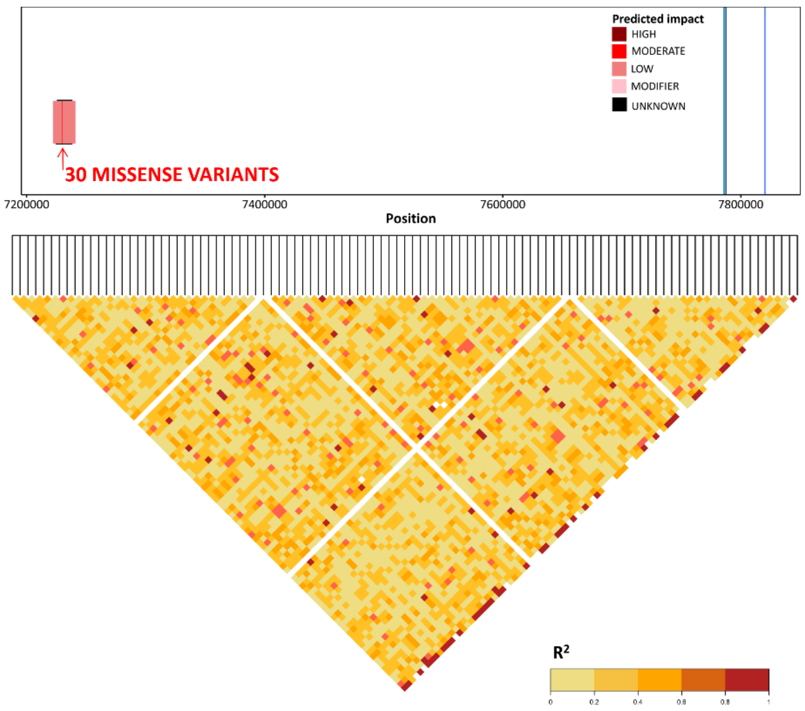


**S21:** Coding region of the gene variants *Bhaga_10.g1460 (ClpS1)*, *Bhaga_10.g1462* (*ASPG1*) and the significant marker on chromosome 10 at ~11.9 Mb in blue and pairwise LD heat map from 11.888957 to 11.918957 Mb.

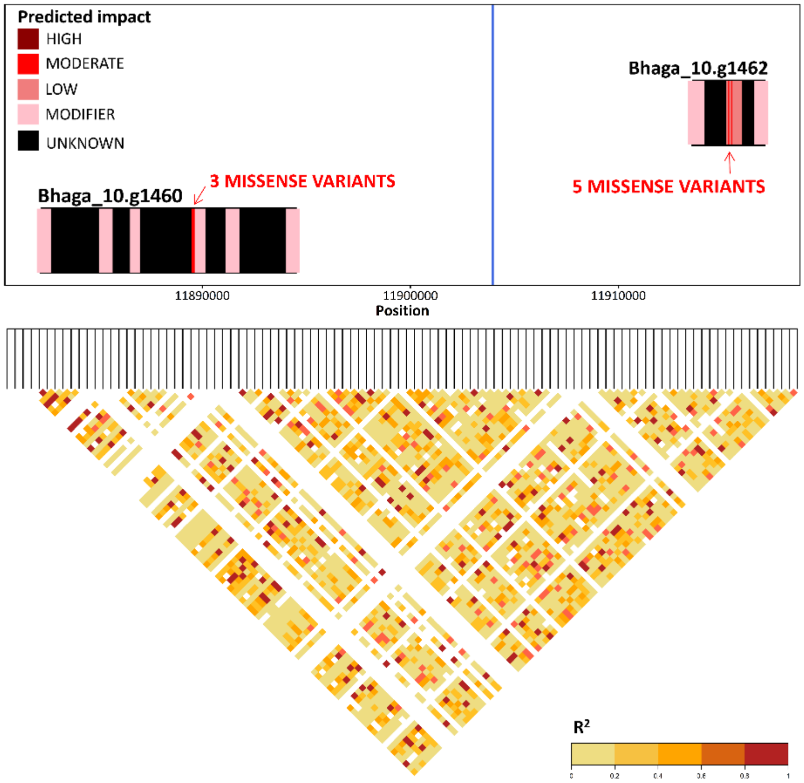

**S22:** Coding region of the gene variant *Bhaga_10.g1651 (BONZAI1)*, *Bhaga_10.g1652* (uncharacterized) and the significant marker on chromosome 10 at ~ 13.44 Mb in blue and pairwise LD heat map from 13420289 to 13440289 Mb.

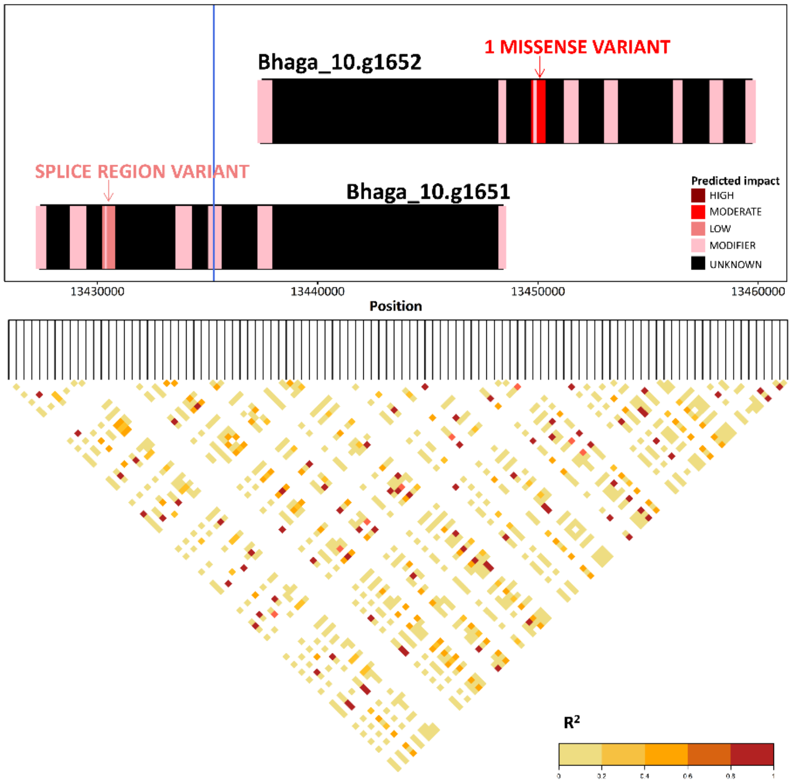


**S23:** Coding region of the gene variant *Bhaga_10.g1543* (*B10-like*) and the significant marker on chromosome 10 at ~ 12.541 Mb in blue and pairwise LD heat map from 12545109 to 12549109 Mb.

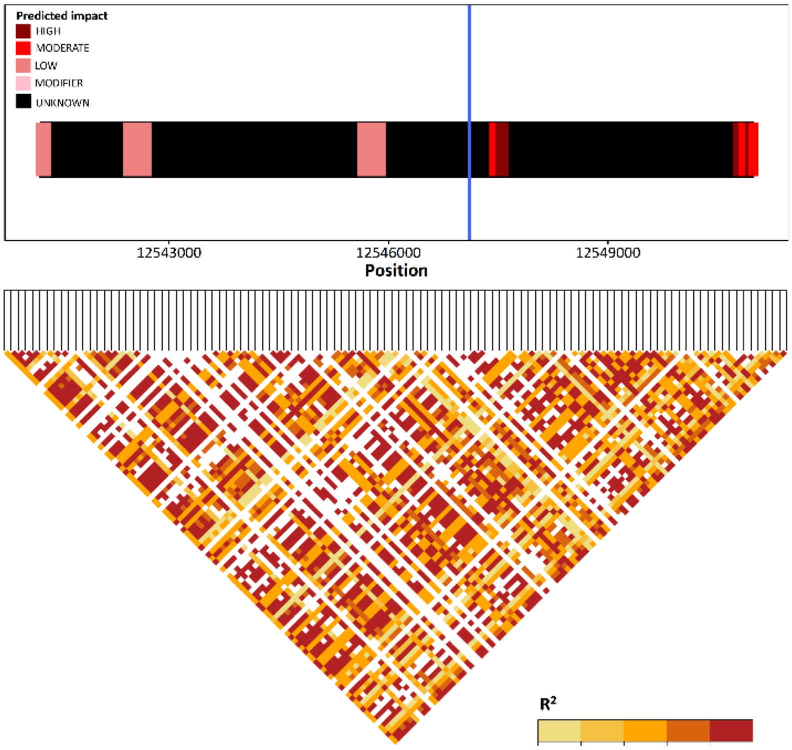


**S24:** List of other genes close to markers associated with stomatal density but without a direct impact on stomatal density based on the literature review

| Gene variant name | Gene | Gene description based on the literature review |
|---|---|---|
| Bhaga_8.g506 | **TFbHLH117** | *TFbHLH117* is a transcription factor of basic-helix-loop helices (bHLH), which is one of the largest families of transcription factors found in plants (Gao and Dubos 2024). |
| Bhaga_10.g752 | **Histone deacetylase HDT1-like** | *Histone deacetylase HDT1-like* appears to be involved in stem vascular development (Zhang et al. 2019b) |
| Bhaga_10.g854 | **Germin-like protein 9-3** | Germin-like protein 9-3 was observed to be involved in a broad-spectrum of disease resistance in rice (Manosalva et al. 2009). Gangadhar et al. (2021) reports that improved heat stress tolerance was observed in potato due to germin-like proteins. |
| Bhaga_10.g867 | **Capsanthin synthase chromoplastic-like** | Capsanthin synthase chromoplastic-like is mainly studied and observed in species of the *Capsicumin* genus like carrot (Deng et al. 2024), paprika, pepper or chili pepper (Piano et al. 2019), but also found in some species of *Lilium leichtinii* (lily) (Valadon and Mummery, 1976) and asparagus (Deli et al. 2000). |
| Bhaga_10.g873 | **mRNA-decapping enzyme-like protein** | The gene mRNA-decapping enzyme-like protein regulates mRNA decapping (Vidya and Duchaine 2022). Decapping of the mRNA means the removal of the m7G cap, which leads quickly to mRNA degradation (Vidya and Duchaine 2022). |
| Bhaga_10.g1543 | **Glycosyltransferase BC10-like** | Glycosyltransferase BC10-like belongs to the family of glycosyl transferases. This gene was only studied in rice (*Oryza sativa* L.) (Zhang et al. 2016) and required for regulation of cellulose biosynthesis in the cell wall (Zhang et al. 2016). |
| Bhaga_10.g1562 | **Kinesin-like protein NACK1** | Kinesin-like protein NACK1, which is a key regulator of plant cytokinesis (Sasabe et al. 2011). Kinesin-like protein NACK1 forms with NPK1 a complex (Nishihama et al. 2002). This complex stimulates kinase activity and plays an essential role in intracellular events that lead to cytokinesis (Nishihama et al. 2002). |
| Bhaga_10.g1674 | **poly(A) polymerase I-like gene** | The poly(A) polymerase I-like gene is responsible 3'polyadenylation which leads to rapid degradation of 3' to 5' exonucleases (Feng and Cohen 2000). This gene was so far only studied in E. coli (Feng and Cohen 2000). |

**S25:** Correlation between stomatal density and observed genotypes at the significant marker ($p \leq 0.000001$) on chromosome 10 at ~7.82 Mb. The coefficient of determination $R^2$ was 0.2353 (p ~0).

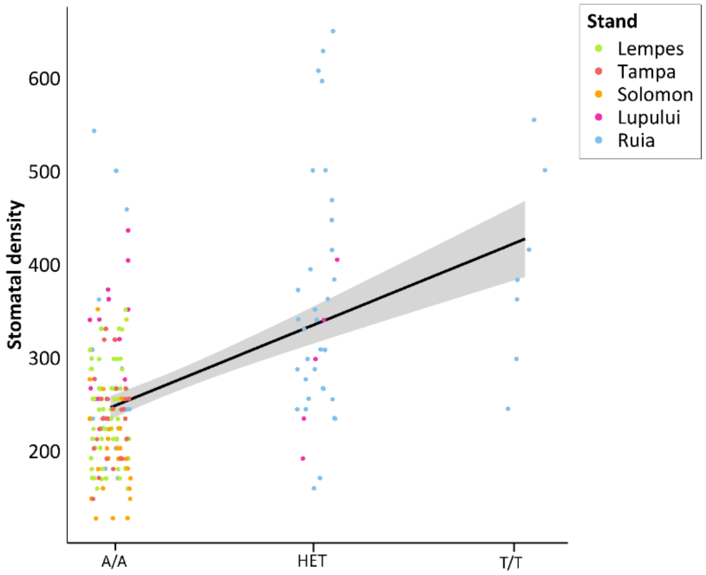
